# Pheno-MYCN maps the morphological footprint of MYCN amplification in paediatric neuroblastoma

**DOI:** 10.64898/2026.08.20.745848

**Authors:** Binghao Chai, Olga Fourkioti, Reed Naidoo, Matt De Vries, Saira Waqar Ahmed, Thomas Jacques, Sally George, Louis Chesler, John Ciaran Hutchinson, Chris Bakal

## Abstract

MYCN amplification and segmental chromosomal abnormalities have long been prognostic markers in paediatric neuroblastoma, and are often accompanied by morphological changes that portend high-risk disease. Yet quantitative morphological assessment is limited to countable features, such as the mitosis-karyorrhexis index. Other high-risk features, such as nuclear size, pleomorphism and anaplasia, lack formal criteria. This leaves a gap: current simple pathology metrics cannot capture, or localise, the full morphological landscape of high-risk biology, such as disease driven by MYCN. Seeking high-risk cases that standard criteria may miss, we developed Pheno-MYCN, a weakly supervised framework linking slide-level MYCN prediction on 189 H&E whole-slide images to interpretable phenotypic sub-populations that expert review mapped to neuroblastoma morphologies. The aim is not a stronger classifier, but to reveal the morphological basis of MYCN prediction. Cell-level profiling found MYCN-associated differences across sub-populations. Amplified tumour was densely cellular but disorganised, with less connected networks, lower cellular diversity and greater abundance of haemorrhagic and acellular necrotic regions. MYCN-amplified-like tissue was identifiable per slide from these features alone (AUC 0.93-1.00, leave-one-slide-out) and traced as a continuous gradient within tumours. Thus MYCN amplification leaves a concrete, interpretable footprint, readable and localisable on routine H&E.

## 1 Introduction

Neuroblastoma is the most common extracranial solid tumour in children and arises from precursor cells of the sympathetic nervous system, typically developing in the adrenal gland or paraspinal regions of the abdomen or chest [1]. Its clinical behaviour is highly heterogeneous, ranging from spontaneously regressing or differentiating tumours to aggressive metastatic disease. Histopathological risk assessment is supported by the International Neuroblastoma Pathology Classification (INPC), which integrates patient age with morphological features such as the mitosis-karyorrhexis index (MKI) to provide prognostic information [2, 3].

Alongside these clinicopathological features, molecular alterations also play a central role in risk stratification. MYCN amplification, located on chromosome 2p24, is one of the strongest adverse prognostic markers in neuroblastoma and is associated with advanced disease, rapid tumour progression, and poor outcome despite intensive treatment [4–7].

Given its prognostic importance, accurate assessment of MYCN status is essential for risk stratification and treatment planning. MYCN amplification is routinely assessed using molecular assays, most commonly fluorescence in situ hybridisation (FISH), which directly visualises MYCN copy number in tumour cells [8]; quantitative PCR (qPCR) can provide complementary information in selected settings [9]. These assays give an accurate readout of copy number, but they can be spatially heterogeneous within a single tumour [10, 11], as such, morphological correlates of MYCN amplification through space (aside from mitosis and apoptosis) remain incompletely understood.

Recent advances in digital pathology and deep learning have made it possible to extract clinically and biologically relevant information from histological images, including subtle tissue patterns, cellular-level information, and morphology associated with diagnosis or prognosis [12–14]. We have developed SurvivMIL, a multimodal multiple-instance learning pipeline that fuses whole-slide images with patient clinical records to stratify survival in high-risk disease and identify poor-outcome patients who might otherwise be overlooked [15]. SurvivMIL, like most computational-pathology models, is built to predict; its measure of success is stratification accuracy. The present study reverses that emphasis: prediction serves only as a probe.

Establishing a direct link between MYCN amplification and H&E morphology would do more than make MYCN status readable from routine slides: since tissue morphology is the visible output of the tumour’s underlying biology, an interpretable morphological signature of MYCN amplification could open avenues to understand how MYCN shapes neuroblastoma tissue. These observations motivate a central question: can weakly supervised analysis of H&E whole-slide images not only detect morphological patterns associated with MYCN amplification in neuroblastoma, but also represent them as an interpretable set of tissue patterns and locate them across the tumour, rather than producing a slide-level prediction alone?

To address this, we introduce Pheno-MYCN, a weakly supervised framework that links MYCN prediction with interpretable phenotypic discovery in H&E whole-slide images (WSIs) of paediatric neuroblastoma. Rather than treating MYCN prediction as a purely slide-level label, Pheno-MYCN also breaks each slide into a small set of recurring, visually distinct tissue patterns (its phenotypic clusters) and measures how much of each a tumour contains. We first benchmark its predictive performance against established weakly supervised WSI classifiers, and describe each slide as a profile of its tissue make-up, the proportion of each pattern it contains that can be compared between MYCN-amplified and non-amplified tumours. We then ask what these recurring tissue patterns represent under the microscope. A specialist paediatric pathologist reviews representative regions from each pattern and links them to recognisable neuroblastoma appearances, including loose and dense tumour, neuropil-poor or more differentiated tumour, haemorrhagic and acellular necrosis, paucicellular tissue, and technical artefact. Using cell-level spatial profiling, we examine what distinguishes MYCN-amplified from non-amplified tissue within each sub-population, and find that MYCN leaves a mark in all of them but through a different kind of feature in each. The tumour regions differ in their cellular architecture: densely packed yet disorganised and less varied; whereas the haemorrhagic and acellular necrotic regions differ mainly in how much of them is present. Finally, we perform an exploratory analysis of patient overall survival to examine whether the H&E-derived tissue patterns or phenotypic clusters, individually and as a joint composition, carry prognostic information beyond MYCN amplification status. Overall, this study asks whether routine H&E morphology contains visible, interpretable traces of MYCN amplification, and how those traces are resolved at the cellular level and organised as a heterogeneous set of patterns across the tumour.

## 2 Results

### 2.1 Pheno-MYCN links MYCN prediction to interpretable phenotypic discovery in H&E whole-slide images

Pheno-MYCN is designed to identify MYCN-associated morphological signals in H&E-stained slides of paediatric neuroblastoma and to expose these signals as interpretable phenotypic patterns. The model operates in a weakly supervised setting, using slide-level MYCN labels without requiring manual regional or cellular annotations. As shown in Fig. 1, Pheno-MYCN combines a discriminative attention-based multiple-instance learning (MIL) branch for slide-level MYCN prediction with an auxiliary Gaussian mixture model (GMM) branch that captures MYCN-associated morphological heterogeneity at the tile level. Both branches share the same tile embeddings produced by the UNI encoder [16]: the MIL branch aggregates tiles into a slide-level representation for classification, whereas the GMM branch assigns each tile to a small set of phenotypic clusters, giving every tile a soft phenotype score (its probability of belonging to each cluster) and a hard cluster label. In effect, one branch decides whether a slide is MYCN-amplified while the other describes the recurring tissue patterns behind that decision, so each prediction comes with an inspectable account of the morphology supporting it.

**Fig. 1.**
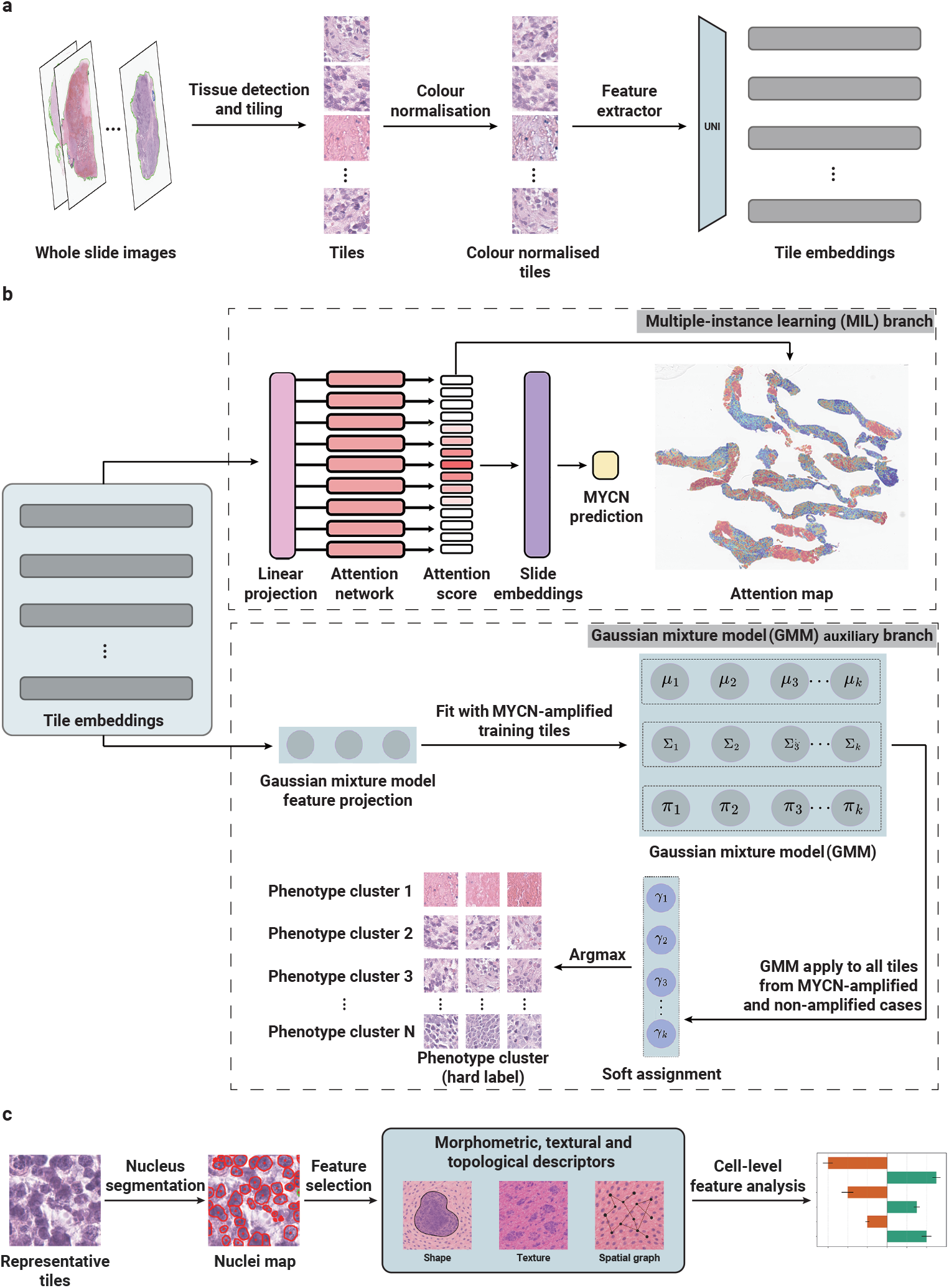
Overview of the Pheno-MYCN framework for MYCN prediction and interpretable phenotypic modelling. **a**, H&E whole-slide images (WSIs) are processed by tissue detection and tiling to generate tissue-containing image tiles. Tiles are colour-normalised and encoded using the UNI feature extractor to produce tile-level embeddings [16]. **b**, Tile embeddings are processed by two complementary branches. The attention-based multiple-instance learning (MIL) branch aggregates tile-level information into a slide-level representation for MYCN amplification prediction and generates attention maps. The auxiliary Gaussian mixture model (GMM) branch is fitted using GMM-projected representations from MYCN-amplified training tiles and then applied to tiles from both MYCN-amplified and non-amplified cases to compute phenotype scores. Hard phenotypic cluster labels are assigned by the maximum phenotype score and used for representative tile retrieval and downstream phenotypic interpretation. **c**, Cell-level analysis workflow. Nuclei within representative tiles from selected GMM phenotypic clusters are segmented to generate nuclei maps, followed by extraction of morphometric, textural, and topological descriptors.

The granularity of the phenotype space, set by the number of clusters the GMM model uses, trades interpretability against resolution, which raises the question of how many clusters best capture the tumours’ morphology. To answer this, we trained Pheno-MYCN with between two and nine GMM phenotypic clusters (*K* = 2 to 9) and chose the number from validation-set performance together with visual inspection of phenotype-cluster coherence in the training data, without using the test set for model selection. Six clusters (*K* = 6) offered the best balance of predictive performance and interpretable, coherent clusters. The resulting cross-validated performance corroborated this choice: *K* = 6 attained a mean test accuracy of 0.889 and F1-score of 0.873, configurations with fewer clusters performed worse (*K <* 6, mean F1 0.766 to 0.813), and additional clusters gave no consistent gain (*K* = 7, *F* 1 = 0.747; *K* = 8, *F* 1 = 0.809; *K* = 9, *F* 1 = 0.789). A four-cluster model (*K* = 4) reached a marginally higher mean AUC (0.917 vs 0.900) but lower accuracy (0.856) and F1-score (0.813), with visibly less coherent clusters, and so was not preferred. We fixed the model at six phenotypic clusters for all subsequent analyses, a setting fine enough to separate distinct tissue phenotypes yet coarse enough to remain interpretable to pathologists (Supplementary Table A2).

To assess whether the interpretable phenotype branch affects predictive accuracy, we compared the six-cluster Pheno-MYCN with two established weakly supervised WSI classifiers that predict MYCN status without any auxiliary phenotype branch: CLAM-SB, an attention-based MIL model [17], and TransMIL, a transformer-based MIL model [18] (Table 1, per-hold-out-split results in Supplementary Table A3). Pheno-MYCN outperformed both baselines on every metric, achieving the highest mean accuracy, F1-score, precision, recall and AUC. These comparisons indicate that Pheno-MYCN can predict MYCN status accurately and group small regions or sub-populations of each slide into recurring tissue patterns (phenotypic clusters), which become the basis for the pathology, cell-level and survival analyses that follow.

**Table 1:** Performance comparison of Pheno-MYCN and baseline weakly supervised WSI classification models. Values are reported as mean *±* SD across ten stratified hold-out splits (per-split results in Supplementary Table A3).

| Model | Accuracy | Precision | Recall | F1-score | AUC |
| --- | --- | --- | --- | --- | --- |
| Pheno-MYCN | $0.89 \pm 0.12$ | $0.89 \pm 0.13$ | $0.87 \pm 0.13$ | $0.87 \pm 0.13$ | $0.90 \pm 0.12$ |
| CLAM-SB | $0.82 \pm 0.09$ | $0.84 \pm 0.19$ | $0.74 \pm 0.14$ | $0.74 \pm 0.16$ | $0.82 \pm 0.11$ |
| TransMIL | $0.82 \pm 0.09$ | $0.81 \pm 0.19$ | $0.76 \pm 0.13$ | $0.76 \pm 0.16$ | $0.84 \pm 0.11$ |

### 2.2 MYCN-amplified and non-amplified tumours differ in morphological sub-populations that map to recognisable neuroblastoma histology

Having fixed the phenotype space at six clusters, we asked whether MYCN-amplified and non-amplified tumours differ in their morphological make-up, that is, in how much of their tissue falls into each phenotypic cluster. The auxiliary Gaussian mixture model (GMM) branch, fitted on tiles from MYCN-amplified training cases, was applied to tiles from both molecular subtypes, so that every tile received a phenotype score for each of the six clusters (the probability that it belongs to that cluster); averaging these scores across a slide summarised each tumour as a profile of phenotypic clusters that could be compared between subtypes across the training, validation and test sets (Fig. 2 **a-c**; per-tile distributions in Supplementary Fig. B1; per-cluster statistics in Supplementary Table A4). Two clusters, 3 and 5, dominated the tissue of both molecular subtypes, together accounting for approximately 91% of the mean slide-level phenotype score in MYCN-amplified cases and 96% in non-amplified cases, while each of the remaining four phenotypic clusters contributed less than 4% on average. Most neuroblastoma tissue therefore falls into two shared phenotypes regardless of MYCN status, with four smaller sub-populations making up the rest.

**Fig. 2.**
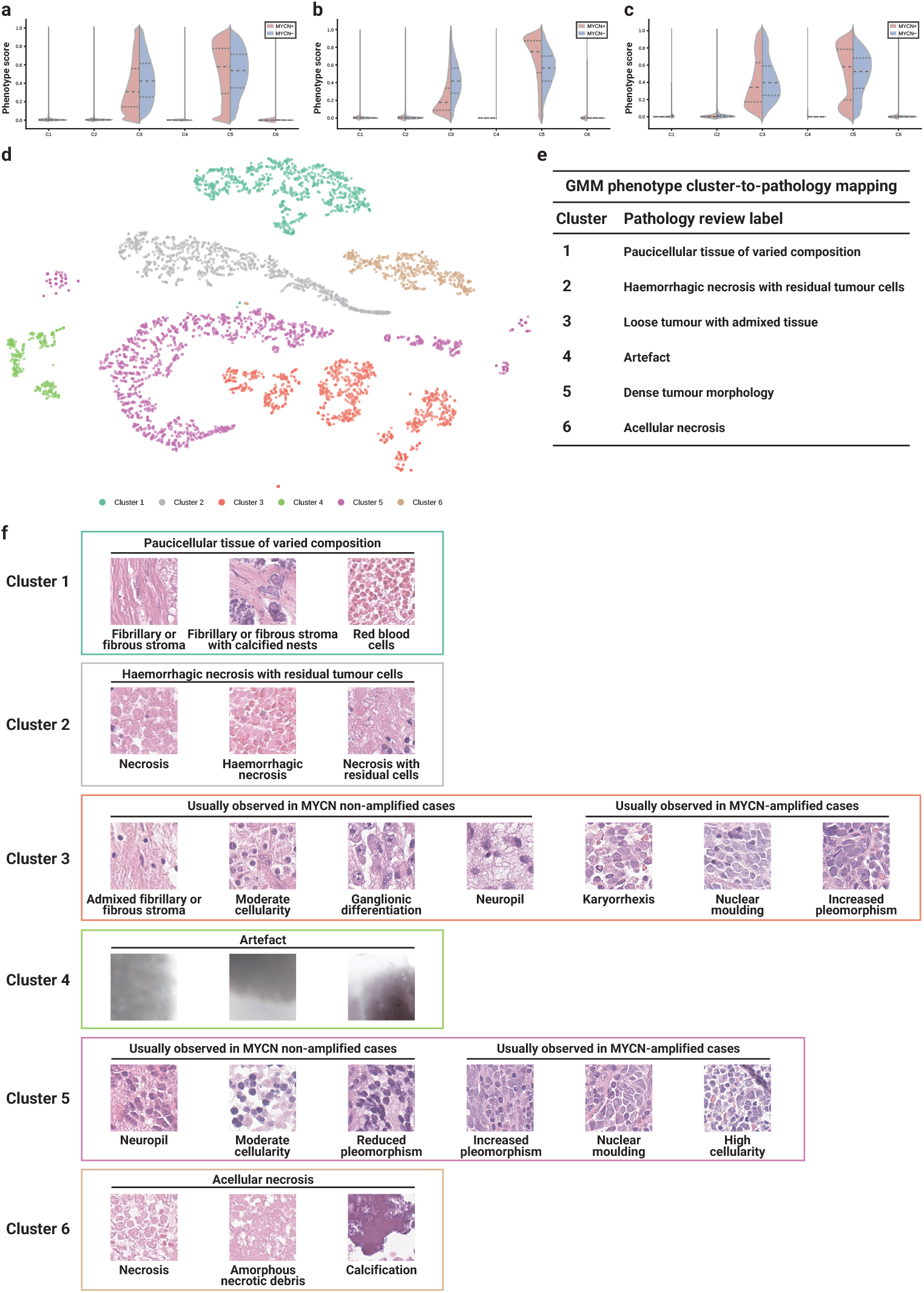
GMM phenotypic clusters differ by MYCN status and map to recognisable neuroblastoma morphologies. **a, b, c**, Violin plots of the phenotype score (the posterior assignment probability) distributions for the training (**a**), validation (**b**), and test (**c**) sets. Each subplot shows the per-tile phenotype score for one of the six phenotypic clusters, stratified by MYCN status; MYCN-amplified and non-amplified cases are shown in red and blue, respectively. The dominant clusters 3 and 5 carry most of the tissue and show the most visible differences in their per-tile score distributions, whereas the minor clusters (1, 2, 4 and 6) are concentrated near zero; the MYCN-associated enrichment of clusters 2 and 6, though small in absolute terms, is quantified in Supplementary Table A4 and described in the main text. **d**, t-SNE embedding of the representative tiles of the six GMM phenotypic clusters (one point per tile), computed from a panel of quantitative per-tile image features and coloured by phenotypic cluster; the six phenotypes occupy largely distinct regions, confirming they are morphologically separable. **e**, Pathology-review label assigned to each learned GMM phenotypic cluster. **f**, Representative tiles for each cluster, reviewed by a specialist paediatric pathologist (J.C.H.). The four clusters capturing tissue other than viable tumour parenchyma (clusters 1, 2, 4 and 6) are each shown as a single set of representative tiles, whereas the two tumour-parenchyma clusters (3 and 5) are shown split by MYCN status. In cluster 3, representative tiles from MYCN non-amplified cases show admixed fibrillary or fibrous stroma, neuropil and ganglionic differentiation with moderate cellularity, whereas tiles from MYCN-amplified cases show karyorrhexis, nuclear moulding and increased pleomorphism; karyorrhexis is scored by the INPC mitosis–karyorrhexis index (MKI). In cluster 5, representative tiles from MYCN non-amplified cases show neuropil-rich areas, moderate cellularity and reduced pleomorphism, whereas tiles from MYCN-amplified cases show increased pleomorphism, nuclear moulding and high cellularity. In both tumour clusters, the MYCN-amplified appearances match the poorly differentiated and, in cluster 3, high-MKI morphology that the INPC classifies as unfavourable histology, whereas the non-amplified tiles are more differentiated and neuropil-rich.

What these six sub-populations represented, and whether any of them distinguished the subtypes, could not be read from the phenotype scores alone. To interpret them, a specialist paediatric pathologist (J.C.H.) reviewed representative tiles (and the whole-slide images they belong to) from every phenotypic cluster, each of which occupied a largely distinct region of image-feature space (Fig. 2 **d**) and showed recognisable patterns (Fig. 2 **e**, **f**). None of the four minor sub-populations corresponded to the viable tumour-cell parenchyma itself: cluster 1 was paucicellular tissue of varied composition (fibrillary or fibrous tissue with calcified nests, fibrin, blood or loose, watery tissue, and a few pale ghost outlines of tumour cells whose nuclei have faded); cluster 4 was technical artefact (out-of-focus regions, tissue folds and air bubbles); and clusters 2 and 6 were haemorrhagic necrosis with residual tumour cells and acellular necrosis, respectively. Of these, the haemorrhagic necrosis (cluster 2) and acellular necrosis (cluster 6) sub-populations showed the clearest difference between the subtypes, both strongly enriched in MYCN-amplified slides (cluster 2: 0.037*±*0.057 versus 0.013*±*0.023, *t* = 4.21, *p <* 0.001; cluster 6: 0.022*±*0.057 versus 0.003*±*0.019, *t* = 3.49, *p <* 0.001; Supplementary Table A4). The sharpest morphological distinction between MYCN-amplified and non-amplified tumours therefore lay not in the bulk of the tissue but in an enrichment of necrotic sub-populations, consistent with the necrosis and vascular disruption that accompany aggressive, MYCN-driven proliferation and that pathologists recognise in aggressive neuroblastoma.

Less apparent was how the two dominant phenotypes, which made up most of the tissue in both subtypes, related to MYCN status. By abundance they barely separated the subtypes: cluster 3 was modestly lower in MYCN-amplified slides (mean 0.387 versus 0.429, *t* = *−*2.01, *p* = 0.046; cluster-3-dominant tiles 29.8% versus 36.3%) and cluster 5 was essentially identical (mean 0.524 versus 0.535, *t* = *−*0.50, *p* = 0.62), yet the per-tile score distributions of both differed in shape between subtypes (two-sample Kolmogorov-Smirnov test with slide-level permutation, cluster 3 *D* = 0.12, *p* = 0.001; cluster 5 *D* = 0.09, *p* = 0.027; Supplementary Table A4). Reviewing their representative tiles split by MYCN status (Fig. 2 **f**), the pathologist identified cluster 3 as loose tumour with admixed tissue and cluster 5 as dense tumour morphology, a more compact tumour phenotype. In both phenotypic clusters, the non-amplified tiles showed neuropil-rich, more differentiated tissue of moderate cellularity, with ganglionic differentiation and admixed fibrillary or fibrous stroma in cluster 3 and reduced pleomorphism in cluster 5; whereas the MYCN-amplified tiles showed increased pleomorphism and nuclear moulding, with high cellularity in cluster 5 and, in cluster 3, prominent karyorrhexis, the apoptotic nuclear fragmentation scored by the MKI. Thus, clusters 3 and 5 did not distinguish MYCN status mainly by how much of the slide they occupied; instead, within these common tumour patterns, MYCN-amplified regions showed a shift towards poorly differentiated and, in cluster 3, high-MKI morphology, mirroring the INPC gradient from favourable to unfavourable histology.

These results show that the learned tissue patterns are both interpretable under the microscope and associated with MYCN status in two ways. First, MYCN-amplified tumours contained more necrotic tissue (haemorrhagic and acellular), small but recognisable components linked to aggressive tumour morphology. Second, the two main tumour patterns were present in both molecular subtypes, but pathologist review showed that MYCN-amplified regions had a more poorly differentiated, high-MKI appearance, with high cellularity, pleomorphism, nuclear moulding and karyorrhexis, whereas non-amplified regions were more neuropil-rich or differentiated. MYCN-associated morphology is therefore not confined to one visible feature or one phenotypic cluster; it appears both as increased necrosis and as a shift in the appearance of the dominant tumour tissue.

### 2.3 What does a tumour’s phenotype composition alone reveal about its MYCN status?

A more stringent test, using no MYCN label at all, was whether that composition would still concentrate MYCN amplification in a coherent morphological subgroup of the cohort. As an unsupervised, slide-level analysis (distinct from the tile-level GMM phenotypes), we grouped all 189 slides by their phenotypic composition alone: the paucicellular tissue (cluster 1) and technical artefact (cluster 4) clusters were excluded, the four tumour-associated clusters (haemorrhagic necrosis, cluster 2; loose tumour, cluster 3; dense tumour, cluster 5; acellular necrosis, cluster 6) were renormalised to sum to one per slide, log-transformed and z-scored, and slides were grouped by Ward’s linkage into five slide-level clusters (Fig. 3). Blind to the MYCN label, the hierarchical clustering separated a minority subgroup of slides (28 of 189; the two necrosis-rich clusters in Fig. 3) that was strongly enriched for MYCN amplification (18 of 28 [64%] versus 31 of 161 [19%]; Fisher’s exact odds ratio 7.6, 95% CI 3.2–18.0, *p* = 3.4 *×* 10*^−^*^6^), whereas the remaining three (Fig. 3) were separated mainly by the relative balance of the loose-(cluster 3) and dense-tumour (cluster 5) phenotypes. On post hoc whole-slide review, the pathologist described the cluster richest in the loose-tumour phenotype as combining the least dense tumour with pale, matrix-rich tissue and, in some slides, features of differentiation. These three MYCN-baseline clusters were each 18–20% MYCN-amplified, so this loose-to-dense balance did not by itself distinguish MYCN status. Even a small sub-population can therefore carry a strong MYCN-associated signal: here the necrotic and haemorrhagic tissue, though a minor fraction of each slide, was enough on its own to separate a MYCN-enriched subgroup of the cohort without the label.

**Fig. 3.**
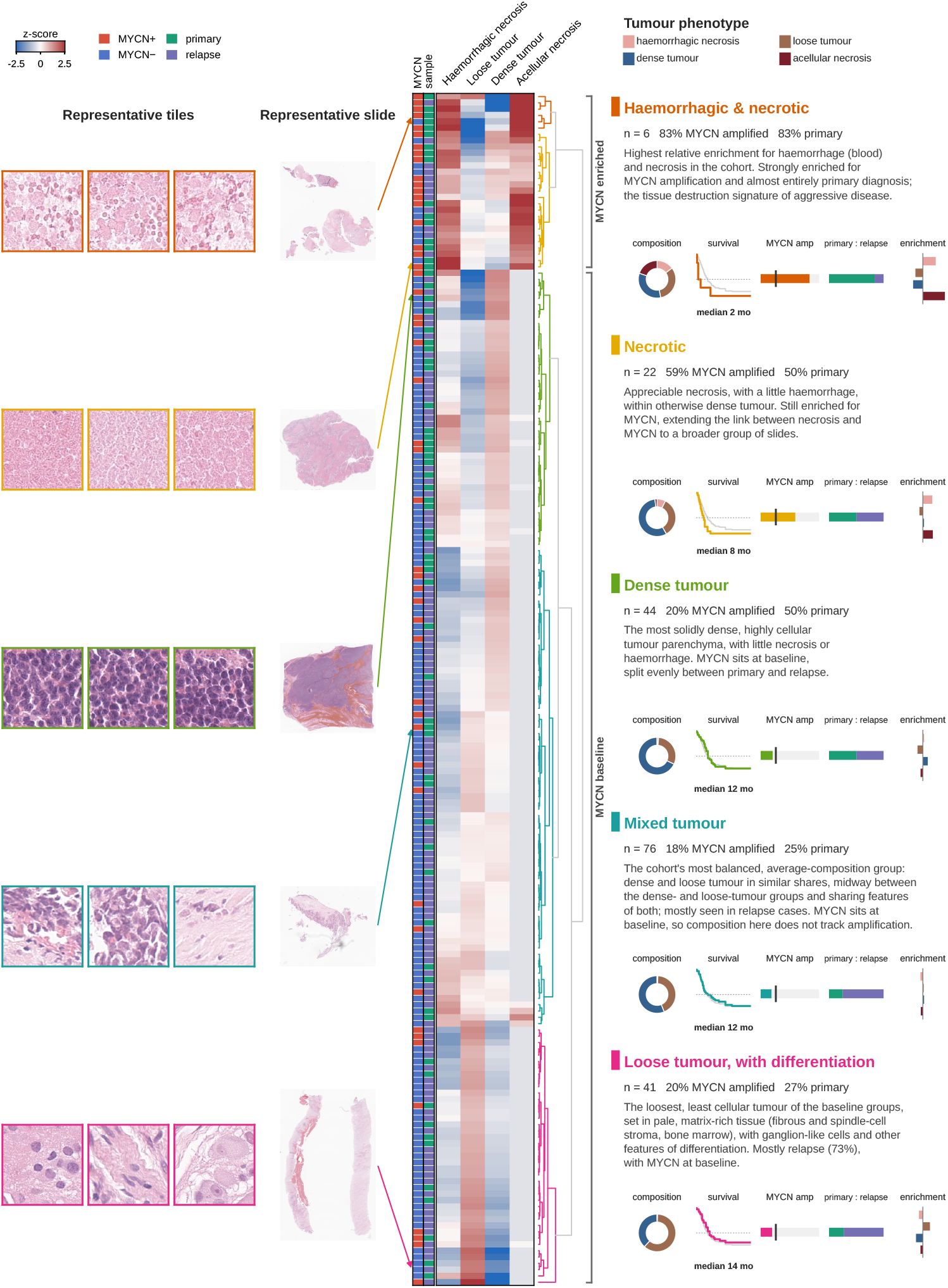
Unsupervised, label-blind slide-level clustering by phenotype composition, with representative morphology and per-cluster clinical and outcome summaries. Hierarchical clustering of all 189 whole-slide images by their GMM phenotype composition, expanded into a per-cluster gallery. For each slide, per-tile phenotypic scores were averaged to a mean score per phenotypic cluster; the paucicellular-tissue cluster (cluster 1) and the artefact cluster (cluster 4) were excluded, and the four tumour-associated phenotypes, haemorrhagic necrosis (cluster 2), loose tumour (cluster 3), dense tumour (cluster 5) and acellular necrosis (cluster 6), were re-normalised to sum to one per slide, log-transformed (log_10_, values floored at 10*^−^*^3^ to avoid amplifying absent components) and z-scored across slides, so that the dominant tumour phenotypes do not swamp the smaller ones. Slides were clustered by Ward’s linkage on Euclidean distances with optimal leaf ordering and the tree was cut into five clusters. Centre, the clustered z-score heatmap (rows, slides; columns, the four tumour-associated phenotypes; red, above the cohort average; blue, below; clipped at *±* 2.5 SD), with colour strips annotating MYCN status (red, amplified; blue, non-amplified) and sample type (green, primary diagnosis; purple, relapse), and the cluster-coloured row dendrogram. The five hierarchical clusters fall into two subgroups: a MYCN-enriched subgroup (haemorrhagic necrotic, necrotic tumours) and three MYCN-baseline clusters (dense tumour; mixed tumour; loose tumour with differentiation, not enriched for amplification. Left, for each cluster, a representative slide and three representative tiles from each, all selected from within the cluster in consultation with a specialist paediatric pathologist (J.C.H.); an arrow links each slide overview to its row in the heatmap. Right, for each cluster, a short description and five summary schematics: (1) composition (donut of the four-phenotype mean composition); (2) survival (Kaplan–Meier per-slide overall survival; cluster in colour, whole cohort in grey; median annotated); (3) MYCN amplification (bar; the tick marks the whole-cohort amplification rate, 26%); (4) primary-to-relapse split; and (5) relative enrichment (mean z-score per phenotype, the signal on which the clustering acts).

The clinical relevance of this subgroup turned on whether it also tracked patient outcome. In descriptive Kaplan-Meier overall-survival curves drawn for each of the five slide-level clusters (Fig. 3), the necrosis-rich clusters enriched for MYCN carried markedly shorter survival than the three MYCN-baseline clusters (median 7.5 versus 12.1 months; log-rank *p* = 0.005). This aligns with established pathology: spontaneous necrosis and haemorrhage mark aggressive, rapidly growing neuroblastoma that outgrows its blood supply, the tissue-level expression of the same high-turnover biology that the INPC grades within the viable tumour through loss of differentiation and a high mitosis-karyorrhexis index (MKI). The comparison is nonetheless exploratory and unadjusted, the per-cluster slide numbers are small, and the enrichment is only partial (36% of the subgroup is non-amplified, and most amplified tumours lie outside it); moreover, the necrotic-cluster scores that characterise the subgroup lose significance after adjustment for MYCN status (Supplementary Table A7). Necrotic burden is therefore best read as a morphological marker of MYCN-driven aggression rather than a prognostic factor in its own right, associated with, but neither necessary nor sufficient for, MYCN amplification and the poor outcome it carries.

These results establish that MYCN amplification is reflected in the morphological composition of neuroblastoma tissue, and that this signal is strong enough to be detected by an analysis blind to the molecular label.

### 2.4 Cell-level profiling shows that MYCN amplification leaves a signature in every sub-population through distinct feature modalities

Having shown that the phenotypic clusters differ by MYCN status and correspond to recognisable histology, we next asked which cellular and spatial features underlie these differences, and whether the same features distinguish MYCN status in every MYCN-associated sub-population. For each of the four MYCN-associated clusters, we selected the ten MYCN-amplified and ten non-amplified slides with the highest mean phenotype score for that cluster and retained every tile the model assigned to it, giving 801 tiles for cluster 2, 1,465 for cluster 3, 1,574 for cluster 5, and 391 for cluster 6. Each tile was characterised by 92 spatial, morphological, and network features derived from HoVer-Net [19] cell segmentation (Fig. 1 **c**). Within each cluster, an L2-regularised logistic-regression classifier trained on these features assigned every tile a soft label, which is its probability of being MYCN-amplified.

The cell-level make-up of a tumour sample alone was enough to detect its MYCN status. Under leave-one-slide-out cross-validation (holding out each slide in turn), the per-slide mean soft label separated MYCN-amplified from non-amplified slides with an area under the ROC curve of 0.93 to 1.00 across the four clusters, whereas individual tiles, which overlap substantially, were only weakly separable (AUC 0.61 to 0.71). In every sub-population, MYCN-amplified slides carried a higher per-slide median soft label than non-amplified slides (cluster 2, 0.57 versus 0.44; cluster 3, 0.60 versus 0.42; cluster 5, 0.56 versus 0.45; cluster 6, 0.56 versus 0.46; Fig. 4 **a**), and the same shift was visible in the full tile-level distributions, widest in cluster 3 (Fig. 4 **b**, Supplementary Fig. B2). The shift held at the level of whole slides: in a principal-component analysis of per-slide mean cell-feature vectors, slides separated by MYCN status along the first or second component in all four clusters, most cleanly in the loose-tumour cluster 3 (Fig. 4 **c-f**). Mapping the cluster-3 tiles into a two-dimensional embedding separated them into two main groups and a small island of 84 outlier tiles (Fig. 4 **g**; cluster 5 at Supplementary Fig. B3). One was sparse and hypocellular, and in most of these tiles too few nuclei could be segmented for the cell features to be measured, so the soft label was uninformative there (median 0.59 in tiles from MYCN-amplified and from non-amplified slides alike). The other was cellular tumour with measurable cell networks, and it was within this group that the soft label carried the signal, with a median of 0.55 in tiles from MYCN-amplified slides versus 0.30 in tiles from non-amplified slides. Tiles of both statuses were interspersed within that group, reflecting intra-tumour heterogeneity: even within one tumour, individual regions span a continuous range of phenotypes.

**Fig. 4.**
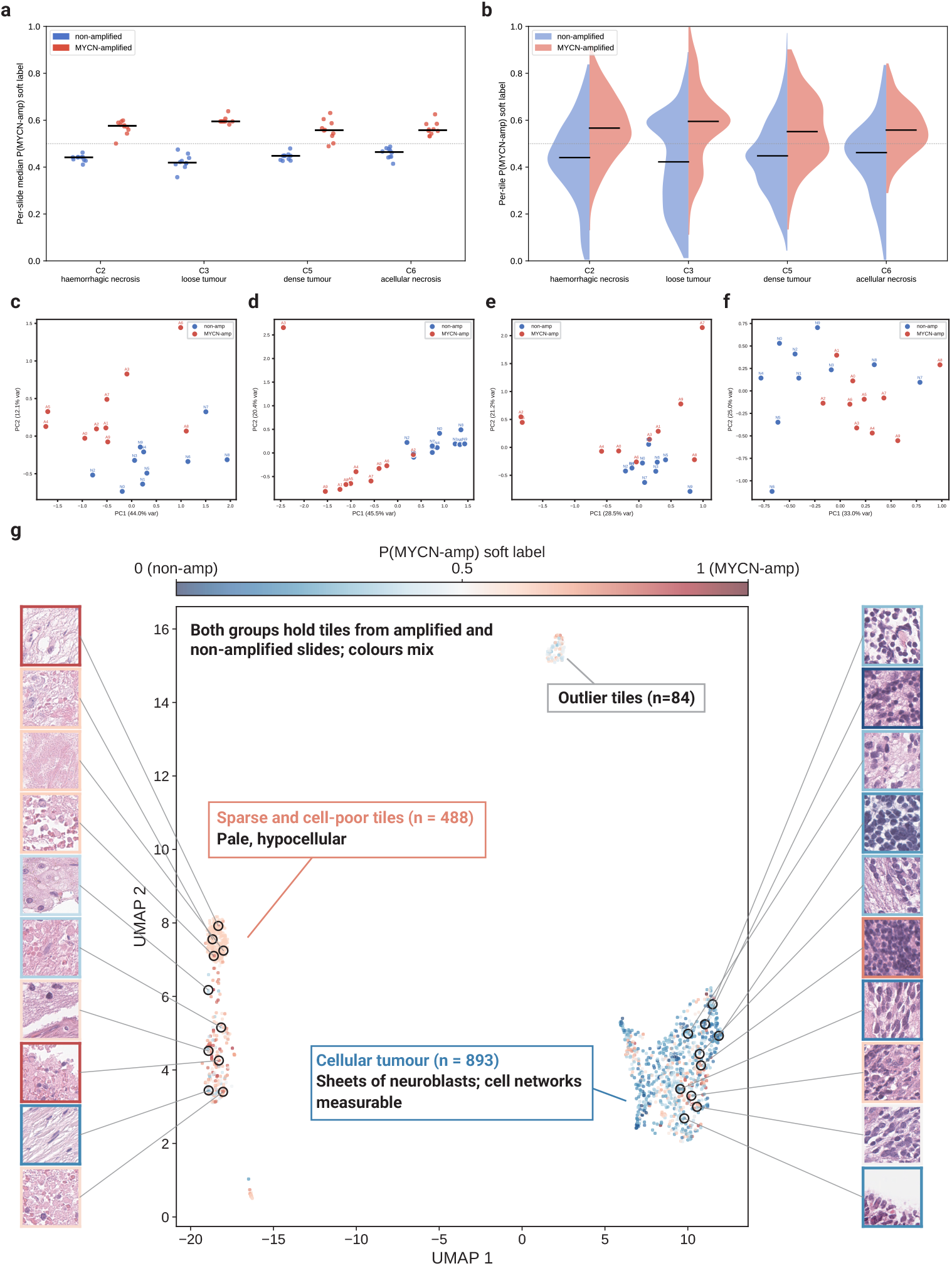
MYCN amplification reshapes the cell-level phenotype of every MYCN-associated sub-population. Latent-space characterisation of four phenotypic clusters using a per-tile soft label: the probability that a tile is MYCN-amplified, from a logistic-regression classifier trained on its segmented-cell features. Non-amplified are shown in blue and MYCN-amplified in red throughout; *n* = 10 non-amplified and 10 MYCN-amplified slides per cluster. **a**, Per-slide median soft label for each cluster (each point, one slide; black bar, group median; dotted line, 0.5). In every sub-population, MYCN-amplified slides carry a higher median soft label than non-amplified slides, the shift being largest in cluster 3. **b**, Split-violin distributions of the pooled per-tile soft labels for each cluster (left/blue, non-amplified; right/red, MYCN-amplified; black bar, median), showing the full tile-level distribution underlying panel a. **c-f**, Principal-component analysis (PCA) of the per-slide mean cell-feature vectors for cluster 2 (**c**), cluster 3 (**d**), cluster 5 (**e**) and cluster 6 (**f**); each point is one slide, labelled by MYCN status, with the variance explained shown on each axis. Slides separate by MYCN status along PC1 or PC2 in every cluster, most cleanly in cluster 3 (with PC1 45.5% of variance). **g**, UMAP embedding of the cluster 3 tiles (one point per tile), coloured by the per-tile soft label. Representative tiles sampled from the left and right of the embedding are shown around it; each tile’s coloured border encodes its own soft label. The embedding separates into two main groups, plus a small island of 84 outlier tiles (top). The smaller group (left) is sparse and hypocellular, and the soft label is uninformative there, sitting near 0.59 irrespective of MYCN status. The larger group (right) is cellular tumour with measurable cell networks, and there the soft label tracks MYCN status (median 0.55 in tiles from amplified versus 0.30 in tiles from non-amplified slides), tiles of both statuses being intermixed within the group. The soft label is a per-tile model prediction rather than the slide’s FISH result, and both groups contain tiles from amplified and from non-amplified slides; distances between groups in a UMAP embedding are not quantitatively interpretable.

To identify which features carried the MYCN signal, we compared every cell-level feature between MYCN-amplified and non-amplified tiles within each cluster using linear mixed-effects (LME) models, with a per-slide random effect to account for the correlation between tiles from the same slide (Fig. 5 **a**; Supplementary Table A5). The discriminating features differed sharply between the tumour and the necrotic clusters. In the loose- and dense-tumour clusters (3 and 5), MYCN-amplified tiles showed coordinated reductions in spatial-network connectivity (closeness and degree centrality, clustering coefficient, and degree) and in cell-population diversity (Simpson, Shannon, and richness indices), together with raised necrotic-cell counts; the effect was broadest and strongest in cluster 3 (31 features significant, 28 of which reduced in MYCN-amplified tiles) and narrower in cluster 5 (11 significant, mainly closeness and degree centrality). MYCN-amplified tumour is classically hypercellular, yet its cell networks were less connected and less diverse. This indicates that amplification is associated with a disrupted, monomorphic tissue architecture, the cellular correlate of karyorrhexis and loss of differentiation, rather than simply denser tissue. The two necrotic clusters (2 and 6) are largely acellular tissue end-states rather than viable cell populations. In both, none of these architectural features reached significance, and in the acellular-necrosis cluster 6 most could not be computed at all because too few nuclei could be segmented. Their only consistent differences were in colour and intensity (hue, redness, and brightness), all raised in MYCN-amplified tiles. We interpret these colour differences cautiously, because they may reflect tissue composition or residual staining and scanning variation rather than a distinct cellular programme. A SHAP analysis of the same classifier independently up-weighted, in the tumour clusters, cellular-diversity indices, neuroblast count, and nuclear size and shape variability (Fig. 5 **b, c**; Supplementary Fig. B4, Supplementary Table A6), broadly consistent with the mixed-effects result. Different sub-populations were therefore distinguished by different feature modalities, the tumour clusters by their disrupted spatial architecture and reduced diversity and the necrotic clusters by colour and intensity. For the latter, the link to MYCN rests chiefly on their greater abundance in amplified tumours, the burden of necrosis and haemorrhage that marks unfavourable neuroblastoma histology.

**Fig. 5:**
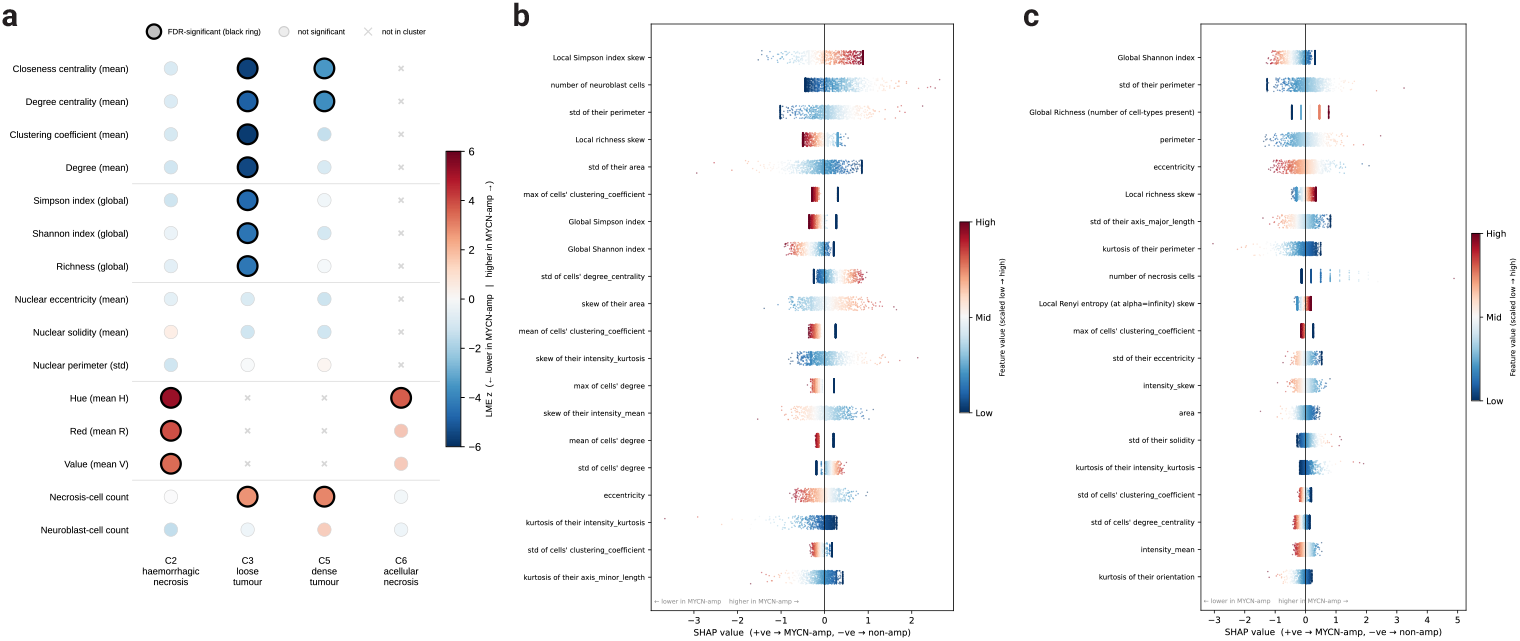
Different histological sub-populations are distinguished by different cell-feature modalities. Cell-level features were aggregated per tile and compared between MYCN-amplified and non-amplified tumours within each of the four MYCN-associated phenotypic clusters (cluster 2, haemorrhagic necrosis; cluster 3, loose tumour; cluster 5, dense tumour; cluster 6, acellular necrosis). **a**, Linear mixed-effects effect map. Each dot is one feature (rows, grouped by modality: spatial connectivity, population diversity, nuclear morphology, colour/texture, cell counts) in one cluster (columns), coloured by the mixed-effects z-statistic for amplification status. A black ring marks effects significant after Benjamini–Hochberg FDR correction (*q <* 0.05); faint dots are tested but non-significant; *×* denotes a feature absent from that cluster’s filtered set. The two tumour clusters (3 and 5) show coordinated reductions in spatial connectivity (and, in cluster 3, population diversity) in MYCN-amplified tumours, together with raised necrotic-cell counts, whereas the two necrotic clusters (clusters 2 and 6) differ only in colour or intensity. Different sub-populations are separated by different feature modalities. **b**, **c**, SHAP attribution for an L2-regularised logistic-regression classifier of tile-level MYCN-amplification status, for the loose-tumour cluster 3 (**b**) and the dense-tumour cluster 5 (**c**). Each point is one tile; the x-axis is the SHAP value (positive values push the classifier towards MYCN-amplified, negative towards non-amplified) and colour encodes the scaled feature value. Features are ordered by mean absolute SHAP value. The classifier independently up-weights cellular-diversity indices, neuroblast and necrotic counts, and nuclear-morphology variability, convergent with the mixed-effects analysis in panel **a**. LME, linear mixed-effects; FDR, false-discovery rate; SHAP, SHapley Additive exPlanations.

Across all four sub-populations, MYCN amplification left a clear cell-level signature, but through different feature modalities in different tissue types. These are two readouts of one biology, that of an aggressive, undifferentiated, necrosis-prone tumour, and they recapitulate at the level of individual cells the same unfavourable-histology features that pathologists assess by eye. The cellular make-up of a tumour sample is therefore not only predictive of MYCN status but interpretable in terms of established histopathology, in every sub-population the model identifies.

### 2.5 Do the learned phenotypes carry prognostic information beyond MYCN status?

To place the learned morphology in a clinical context, we asked whether it carries prognostic information and whether it complements the molecular assay, in an exploratory slide-level survival analysis. Of the 189 slides in the model cohort, 187 had usable overall-survival data; two slides from one patient were excluded because the recorded date of death preceded the resection date. As expected, MYCN amplification was strongly prognostic in this cohort; Kaplan–Meier overall survival stratified by per-slide FISH MYCN status separated cleanly throughout follow-up (median 14.7 months in non-amplified versus 5.4 months in MYCN-amplified slides; log-rank *p <* 0.001; Fig. 6 **a**). This recovered the established adverse prognosis of MYCN amplification and set the benchmark against which any morphology-derived survival signal had to be measured.

**Fig. 6.**
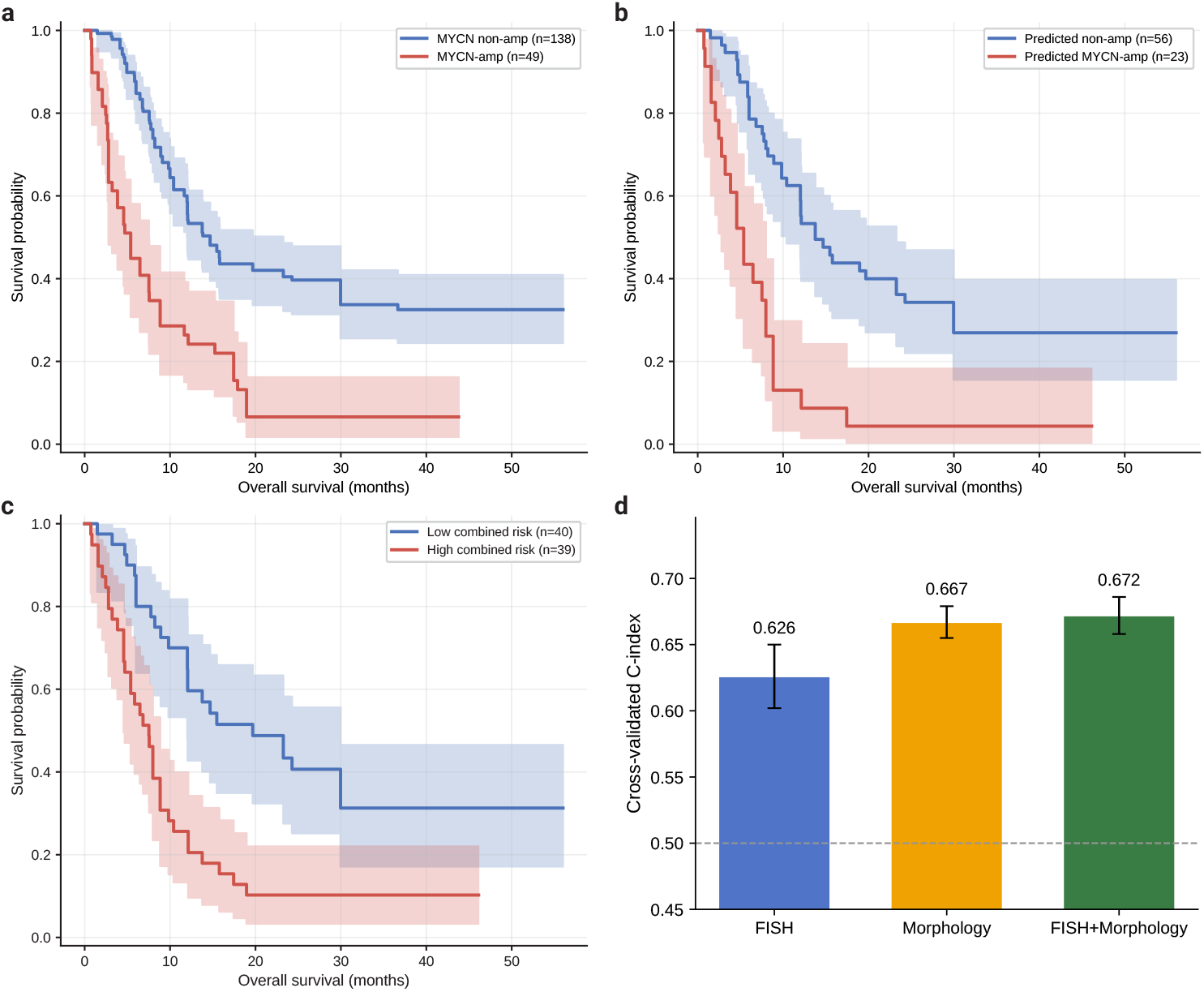
H&E morphology recovers MYCN’s prognostic value and adds to it. Kaplan–Meier overall-survival curves and Cox proportional-hazards models for the slide-level survival cohort (*n* = 187 slides). **a**, Overall survival by FISH MYCN status across the full cohort (138 non-amplified, 49 amplified). MYCN amplification is strongly adverse, with a median overall survival of 14.7 versus 5.4 months (non-amplified versus amplified; log-rank *p <* 0.001). **b**, Overall survival by the H&E-derived, out-of-fold predicted MYCN call, on the 79 held-out slides for which a prediction was available (56 predicted non-amplified, 23 predicted amplified). The morphology-based call recovers the same prognostic separation, with median overall survival of 13.8 versus 5.4 months (log-rank *p <* 0.001). On a continuous scale, the predicted probability of MYCN amplification reaches a Cox concordance index of 0.69 (hazard ratio 1.81 per standard deviation, *p <* 0.001), comparable to FISH MYCN status on the same slides (concordance index 0.65). **c**, Overall survival by a combined MYCN-plus-morphology risk score on the 79 held-out slides. The risk score is the out-of-fold linear predictor of a Cox model combining FISH MYCN status with the continuous H&E-derived probability of MYCN amplification, split at the cohort median (low, *n* = 40; high, *n* = 39). Higher combined risk marks markedly shorter survival, with a median overall survival of 19.7 months in the low-risk versus 7.5 months in the high-risk group (log-rank *p <* 0.001). **d**, Cross-validated discrimination (Harrell’s concordance index) of three Cox models on the same 79 slides: FISH MYCN status alone, the H&E-derived morphology score alone, and the two combined. Bars show the mean concordance index over 20 repeats of 5-fold cross-validation, error bars the standard deviation, and the dashed line marks chance (0.5). The morphology score alone discriminates outcome better than binary FISH (0.667 versus 0.626) and adding it to FISH gives the most prognostic model (0.672; likelihood-ratio test of the morphology score on top of FISH, *p* = 0.027), showing that morphology and MYCN are complementary and most prognostic together. In the Kaplan–Meier panels (**a**–**c**), shaded bands are 95% confidence intervals.

Before turning to the individual phenotypic clusters, we tested whether the morphology as a whole could reproduce this prognostic separation. Using the model’s slide-level MYCN predictions out of fold, so that each slide was scored only by stratified splits in which it had been held out (79 of the 187 slides received such a prediction), we stratified survival by the H&E-derived MYCN call. The morphology-based call recovered essentially the same prognostic split as the molecular assay (median 13.8 months in predicted-non-amplified versus 5.4 months in predicted-amplified slides; log-rank *p <* 0.001; Fig. 6 **b**), and it agreed with FISH on 69 of the 79 slides. On a continuous scale, the predicted probability of amplification discriminated outcome as well as the assay itself (Cox concordance index 0.69, versus 0.65 for binary FISH on the same slides; hazard ratio 1.81 per standard deviation, *p <* 0.001). The morphology that Pheno-MYCN reads is therefore prognostically informative to a degree comparable with the molecular test, which raises the question of whether any of that prognostic value comes from morphology beyond what MYCN status already encodes.

Whether the morphology adds to the molecular assay, or merely mirrors it, is a separate question. On the 79 held-out slides, we compared Cox proportional-hazards models built on FISH MYCN status, on the model’s continuous H&E-derived MYCN score, and on the two combined, summarising each by its cross-validated concordance index over 20 repeats of five-fold cross-validation, and we stratified survival by a combined MYCN-plus-morphology risk score split at the cohort median. The morphology score alone discriminated outcome better than binary FISH (cross-validated concordance index 0.67 versus 0.63) and added to it significantly (likelihood-ratio test *p* = 0.027), so that the combined model gave the highest discrimination (Fig. 6 **d**). The combined risk score separated survival sharply, with a median overall survival of 19.7 months in the low-risk versus 7.5 months in the high-risk group (log-rank *p <* 0.001; Fig. 6 **c**). The morphology that Pheno-MYCN reads therefore carries prognostic information beyond the binary molecular call, adding to the assay rather than merely reproducing it, so that morphology and MYCN are best used together.

A more stringent test is whether the phenotypic clusters themselves carry that added value, taken jointly rather than one at a time. We entered the four tumour-associated cluster scores together as a compositional profile, using the same log transform as the slide-level clustering, in a Cox model evaluated out of fold against MYCN status. The joint composition carried no reproducible prognostic signal (cross-validated concordance index 0.51, at chance) and added nothing to MYCN status (0.59 combined versus 0.61 for MYCN alone; likelihood-ratio test *p* = 0.49), and no single cluster remained significant once MYCN was accounted for (adjusted *p >* 0.11 for every cluster; Supplementary Table A7). The prognostic signal therefore lies in the model’s rich, learned morphological representation rather than in the coarse phenotype-cluster proportions, which are best read as an interpretable description of MYCN-associated tissue rather than as a prognostic index in their own right.

## 3 Discussion

In this study, we introduced Pheno-MYCN, a weakly supervised digital pathology framework designed to link slide-level MYCN prediction with interpretable phenotypic discovery in H&E-stained whole-slide images of paediatric neuroblastoma. Rather than treating MYCN amplification as a purely black-box classification target, Pheno-MYCN combines an attention-based multiple-instance learning classifier with an auxiliary Gaussian mixture model phenotype branch. This design enabled the model to both predict MYCN status and assign tiles to a learned phenotype space that could be inspected using phenotype scores, pathology review, and cell-level spatial profiling. Across the analysed cohort, Pheno-MYCN showed stronger predictive performance than representative weakly supervised baselines, while the learned GMM phenotypic clusters revealed subtype-associated morphological patterns linked to MYCN amplification. This performance matters chiefly as evidence that the learned representation tracks MYCN biology; the contribution lies in what that representation reveals and in the fact that it can be read.

A central contribution of this work is the use of GMM phenotype scores as an interpretable representation of morphological heterogeneity, one that turns a slide-level MYCN prediction into a mechanistic, cellular-level reading of the underlying tissue biology. Rather than identifying a single dominant MYCN-associated pattern, the learned phenotype space captured differential enrichment across multiple phenotypic clusters, including shared tissue contexts as well as tumour-rich morphologies with subtype-associated differences. Expert pathology review mapped these phenotypic clusters to recognisable neuroblastoma features, including cellularity, nuclear moulding, karyorrhexis, neuropil, ganglionic differentiation, necrosis, haemorrhage, and artefact. The model recovered these axes with no morphological supervision, having been trained only on slide-level MYCN labels, so their agreement with the features pathologists already assess (several of which are formal INPC criteria) is independent evidence that the phenotype space reflects clinically meaningful tissue rather than arbitrary image texture.

Cell-level spatial profiling further linked selected MYCN-associated clusters to quantitative features of nuclear organisation, cellular packing, necrosis-associated morphology, and neuroblast morphometry, providing an inspectable layer between slide-level molecular prediction and the histology itself.

These morphological patterns can be read as the histological footprint of MYCN itself. This footprint is not a single appearance but a heterogeneous mix of morphologies, from the disrupted architecture of the tumour parenchyma to the necrosis and haemorrhage that accompany it. MYCN is a transcription factor that sustains proliferation and holds neuroblasts in an undifferentiated state, and the MYCN-amplified phenotype that the model surfaced is the visible expression of that programme: high cellularity, nuclear moulding, abundant karyorrhexis, and a loss of the neuropil and ganglionic differentiation seen in more favourable tumours. The karyorrhexis is informative in its own right, because it reflects the high cell turnover that MYCN drives by pushing proliferation while also priming cells for death. At the cellular level, MYCN-amplified tumour regions showed reduced spatial connectivity and, most clearly in the loose-tumour cluster, lower population diversity, the signature of a monomorphic sheet of poorly differentiated neuroblasts rather than organised, architecturally diverse tissue. The necrosis and haemorrhage enriched in MYCN-amplified tumours fit the same picture, arising when rapid and disorganised growth outstrips and damages the blood supply. In summary, these features are not independent markers but downstream consequences of a single upstream driver, consistent with the survival analysis.

We then asked, in an exploratory analysis, what the learned morphology contributes to patient outcome. The model’s H&E-derived MYCN prediction, generated without any molecular input, recovered almost exactly the prognostic separation that the FISH assay provides (Fig. 6 **b**) and added to it rather than merely reproducing it, being most discriminating when combined with the molecular call (Fig. 6 **c**, **d**). This prognostic value was carried by the model’s rich learned representation rather than by the interpretable phenotypic clusters, which, entered jointly as a composition, added nothing beyond MYCN status. The interpretable clusters are therefore best read as a description of MYCN-associated biology rather than as independent prognostic markers, tracking the same aggressive, undifferentiated, necrosis- and haemorrhage-prone phenotype that MYCN drives and that connects the cell-level signatures to the unfavourable histology pathologists already grade. Read this way, Pheno-MYCN both characterises MYCN-associated morphology and complements the molecular assay for outcome, its prognostic value lying in the combination rather than in any single phenotypic cluster acting on its own.

Clinically, MYCN amplification remains a molecular biomarker that should continue to be assessed using established assays such as FISH, and Pheno-MYCN is therefore not intended to replace molecular testing. Its value is complementary, and it points to two practical uses. First, the H&E-derived MYCN call recovers the prognostic stratification of the assay, so it could act as a low-cost triage that flags cases for confirmatory FISH. This is most useful where molecular testing is delayed, limited, or unavailable, such as centres without routine molecular diagnostics. Second, MYCN amplification can be focal, and a single sampled block may miss it [10, 11], so the model’s tile-level per-region map of MYCN-associated morphology could point to the areas most likely to be amplified and direct molecular testing toward them. More broadly, the same principles may extend beyond neuroblastoma. Other diagnostic settings rely on molecular assays to distinguish morphologically overlapping entities, such as MDM2 testing in the differential diagnosis of lipoma and atypical lipomatous tumour [20], or EWSR1 rearrangement testing in Ewing sarcoma and related small round cell tumours [21]. Weakly supervised phenotypic modelling may therefore offer a general strategy for asking whether routine H&E images contain interpretable correlates of molecular alterations, including copy-number changes, rearrangements, fusions, and other genomic and transcriptomic states.

This work has several limitations. Neuroblastoma is a rare paediatric tumour, so the cohort was modest and single-centre, with relatively few MYCN-amplified cases, and the survival analysis was exploratory, using the H&E-based MYCN prediction only on the slides held out during cross-validation. Most samples were taken at relapse, so some morphology may reflect prior treatment, such as therapy-induced necrosis, fibrosis or maturation, rather than MYCN biology alone. Splitting and evaluation were performed at the slide level rather than the patient level, because MYCN amplification is a property of the individual tumour sample and a few patients contributed both amplified and non-amplified slides. We found no evidence that this inflated performance, as accuracy on test slides whose patient was absent from training was no lower than overall, but larger multi-institutional cohorts will be needed to confirm generalisability across centres, scanners, staining protocols, and patient populations. Indeed, a recent study of soft-tissue tumour classification found that classifiers built on pathology foundation models can lose accuracy when tested on slides stained at other institutions or digitised on different scanners, while remaining markedly more robust than conventional convolutional neural networks [22]. In addition, MYCN status was treated as a binary FISH label, and the framework does not model copy-number burden, co-occurring molecular alterations, or borderline states. Finally, the per-tile soft label measures morphological similarity to MYCN-amplified tissue rather than a measured per-region genotype, and in the two necrotic clusters the cell-level signal is carried largely by colour and intensity, features of aggressive tissue that are not specific to MYCN.

Future studies should investigate how phenotype-discovery models such as Pheno-MYCN behave across broader molecular and clinical contexts. External validation will be essential to determine whether the learned MYCN-associated phenotypic clusters are stable across independent cohorts and whether similar phenotype scores can be recovered when models are trained on data from different institutions. One observation deserves particular attention. A small number of slides were discordant between the H&E call and FISH, and those predicted amplified by morphology but non-amplified by FISH tended toward shorter survival. The numbers are very small and only hypothesis-generating, but if the pattern holds in a larger cohort it could mean the model is detecting an aggressive, MYCN-like morphology driven by other high-risk mechanisms, such as MYCN over-expression without amplification, 11q deletion, ALK activation, or telomere-maintenance pathways, pointing toward a morphology-based high-risk signal that does not depend on MYCN amplification and that could help identify poor-outcome patients who might otherwise be overlooked. Methodologically, the framework could be extended to multi-task settings that jointly model several molecular alterations, histopathological features, and clinical outcomes, and a natural test would enrich for MYCN-non-amplified high-risk disease in a larger, ideally multicentre cohort. By combining slide-level prediction with tile-level phenotypic modelling and cell-level interpretation, Pheno-MYCN illustrates how computational pathology can move beyond classification alone toward interpretable discovery of molecularly associated tumour phenotypes.

## 4 Methods

### Ethical approval declarations

Tumour samples and linked clinical data were collected under the Stratified Medicine Paediatrics (SMPaeds) and Stratified Medicine Paediatrics 2 (SMPaeds2) programmes, for which The Institute of Cancer Research is the study sponsor. SMPaeds was approved under ethical approval CCR4958 (IRAS 246557) and SMPaeds2 under ethical approval CCR6062 (IRAS 346034). Archival haematoxylin and eosin sections and the corresponding formalin-fixed paraffin-embedded blocks were obtained from the Great Ormond Street Hospital biobank.

### 4.1 Dataset

We employed 189 H&E-stained whole-slide images (WSIs) from 86 patients with high-risk paediatric neuroblastoma. These WSIs were collected and digitised from Great Ormond Street Hospital (GOSH); the Formalin-Fixed Paraffin-Embedded (FFPE) tissue samples were collected between 2019 and 2023, and the samples were enrolled in the Stratified Medicine Paediatrics (SMPaeds) programme. At the slide level, there are 49 MYCN-amplified (26%) and 140 non-amplified (74%) WSIs; 68 slides were taken at primary diagnosis and 121 at relapse. Patients had a median age at biopsy of 5.7 years (interquartile range 3.8 to 8.6; range 0.9 to 36.0); 44 were female and 42 male. At last follow-up, 60 patients had died and 26 were alive. Patients contributed a median of two slides each (range 1 to 6); 64 of 86 patients had more than one slide, sampling different timepoints (primary and relapse) and sites, and 6 patients had both MYCN-amplified and non-amplified slides (Supplementary Table A1). MYCN status was determined from fluorescence in situ hybridisation (FISH) results and treated as a binary variable, categorised as amplified or non-amplified. During data review, a small number of initially mislabelled cases were identified and corrected in consultation with a pathologist. All cases included in the final analysis, therefore, had confirmed MYCN status, with no borderline, failed, or unknown MYCN labels. Available clinical metadata included patient age, MYCN status, biopsy or resection dates, and outcome-related variables where available.

Because MYCN amplification status is a property of the individual tumour sample rather than of the patient, the slide was used as the unit for splitting. A patient’s slides are taken at different times (primary diagnosis and relapse) and from different anatomical sites, and these samples can carry different MYCN status, with 6 of the 86 patients having both MYCN-amplified and non-amplified slides. Patient identity therefore does not determine the slide-level label, and slides from a given patient could fall in different hold-out splits. Model development used ten repeated stratified hold-out splits, each comprising 171 training, 9 validation, and 9 held-out test slides. An out-of-stratified-split MYCN prediction for a given slide can come only from a split in which that slide was held out for testing. Across the ten splits, 79 of the 189 slides were assigned to a held-out test set in at least one split and so received such an out-of-stratified-split prediction, whereas the remaining slides were used only for training or validation. The H&E-derived MYCN survival analysis (Fig. 6 **b**) was therefore restricted to these 79 slides. For survival analyses, two slides from a single patient were subsequently excluded because the recorded date of death preceded the resection date, leaving 187 slides.

### 4.2 WSI preprocessing and tile-level feature extraction

Whole-slide images were preprocessed using CLAM [17] to generate tissue masks and extract image tiles from tissue-containing regions. Each WSI was tiled into non-overlapping 224 *×* 224 pixel tiles at 40*×* magnification, corresponding to a resolution of 0.252 *µ*m per pixel. To further remove non-informative background areas, pixels with an average intensity above 230 were classified as background. Tiles with less than 60% tissue content were discarded.

To reduce staining variability across samples, stain normalisation was applied using the Reinhard method [23]. The target stain distribution was estimated from the average colour-channel statistics of 100 randomly selected tiles from the dataset. Tissue-containing tiles were then encoded using the UNI feature extractor [16], a ViT-L/16 model based on DINOv2 [24]. These tile-level embeddings were used as input to Pheno-MYCN, a weakly supervised dual-branch framework comprising an attention-based multiple-instance learning classifier for slide-level MYCN prediction and an auxiliary Gaussian mixture model branch for phenotypic modelling.

### 4.3 Attention-based multiple-instance learning (MIL) classifier

To generate slide-level MYCN predictions from tile-level embeddings, we used an attention-based multiple-instance learning (MIL) classifier [25]. For each WSI, the retained tissue tiles were treated as a bag of feature vectors *{****t****_i_}_i=_*_1_*^N^*. Attention weights were computed using the gated attention mechanism. The classifier assigns an attention weight *a_i_* to each tile and aggregates the transformed tile representations ***m****_i_* into a slide-level representation:

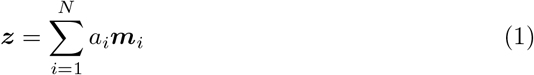

The resulting slide-level feature vector ***z*** was passed to a linear classification layer to predict MYCN amplification status. This classification branch was trained using binary cross-entropy loss with slide-level MYCN labels. The learned attention scores were also used to visualise spatial regions contributing to the slide-level prediction.

### 4.4 Gaussian mixture model phenotype branch

In addition to the attention-based multiple-instance learning branch, Pheno-MYCN includes an auxiliary Gaussian mixture model (GMM) branch designed to model MYCN-associated morphological heterogeneity. Before GMM modelling, tile embeddings were passed through a GMM-specific linear projection layer. The resulting projected tile-level representations were used for GMM fitting, responsibility estimation, and hard component assignment. The fitted GMM defines a set of learned phenotypic clusters that can be used to compare phenotypic similarity and heterogeneity between MYCN-amplified and non-amplified cases.

Let *T*_amp_ = *{****t****_i_}_i=_*_1_*^Namp^* denote the projected tile-level representations extracted from MYCN-amplified training slides. These projected representations were modelled using a GMM with *K* components, where each component *k* is defined by a mixture weight *π_k_*, mean vector ***µ****_k_*, and covariance matrix **Σ***_k_*:

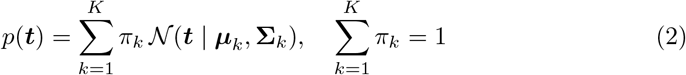

GMM parameters were initialised using a two-step prototype-based clustering strategy inspired by [26]. First, projected tile representations within each WSI were grouped according to their cosine similarity to a slide-level prototype computed from the mean projected tile embedding. These initial groups were then refined across MYCN-amplified slides using an iterative centroid-based clustering procedure to improve within-component coherence. The resulting centroids, empirical covariances, and component proportions were used to initialise ***µ****_k_*, **Σ***_k_*, and *π_k_*, respectively.

During training, the GMM loss was applied only to projected tile representations from MYCN-amplified slides, with *λ* = 0.001 used to weight the GMM negative log-likelihood term in the overall objective:

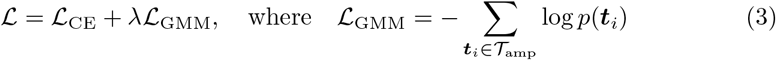

After fitting, the GMM was applied to projected tile representations from both MYCN-amplified and non-amplified slides. Fitting the GMM on MYCN-amplified tiles alone defines an intentionally MYCN-associated phenotype space: the phenotypic clusters capture the morphological patterns characteristic of MYCN-amplified tumours, and applying them to non-amplified tiles describes non-amplified tissue relative to this MYCN-amplified reference, by design. For each projected tile representation, the posterior probability, or responsibility, for component *k* was computed as:

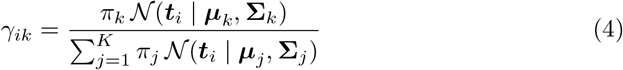

Responsibilities were used as soft component assignments, and hard component labels were obtained by assigning each tile to the component with the highest responsibility. We refer to these GMM components as phenotypic clusters and to the posterior responsibility as the phenotype score.

### 4.5 Model training and baseline evaluation

We trained and evaluated Pheno-MYCN using ten repeated stratified hold-out splits, each comprising 171 training, 9 validation, and 9 test slides, stratified by slide-level MYCN status, with the slide as the unit of splitting and evaluation. MYCN amplification is a property of the individual tumour sample rather than of the patient: 6 of 86 patients contributed both amplified and non-amplified slides, sampled at different timepoints (primary diagnosis and relapse) and sites, so patient identity does not determine the slide-level label. Given the limited size of this rare-tumour cohort, we deliberately used a large training fraction, retaining the small held-out validation and test sets for model selection and evaluation. Because each split’s test set is small, the individual estimates are noisy, so performance is reported as the mean and standard deviation across the ten splits. Slides from a given patient could fall in different splits, so to check for patient-specific shortcuts, we recomputed test performance on the subset of test slides whose patient was absent from the corresponding training split. Accuracy and discrimination on these fully held-out patients were no lower than the overall test performance, indicating that the reported metrics are not driven by patient-level leakage. The attention-based classification branch was trained using slide-level MYCN labels from all training cases, whereas the GMM loss was applied only to GMM-projected tile representations from MYCN-amplified slides.

The entire model was optimised using the Adam optimiser [27] with a learning rate of 0.0002 and a weight decay of 10*^−^*^5^. Training was conducted for up to 50 epochs, with early stopping applied using a patience of 20 epochs. The model with the lowest validation loss was retained for evaluation. Model performance was assessed using accuracy, precision, recall, F1-score, and area under the receiver operating characteristic curve (AUC). For baseline comparison, CLAM-SB [17] and TransMIL [18] were trained and evaluated using the same preprocessed tiles, UNI feature embeddings, hold-out splits, and evaluation metrics as Pheno-MYCN.

### 4.6 GMM phenotypic cluster selection and phenotype score

To select the granularity of the GMM phenotype space, we evaluated Pheno-MYCN configurations with *K* = 2 to 9 GMM phenotypic clusters, assessing each across all ten stratified hold-out splits (Supplementary Table A2). The number of phenotypic clusters was chosen on the basis of validation-set predictive performance together with visual inspection of cluster coherence on the training data; the held-out test set was not used for this choice. On this basis, a six-cluster configuration was fixed before any downstream phenotypic analysis (phenotype scoring, pathology review, and cell-level analysis).

Using the fixed six-cluster configuration, GMM phenotype scores were computed for GMM-projected tile representations from MYCN-amplified and non-amplified slides. For each slide, per-tile phenotype scores were averaged across all tiles for each GMM phenotypic cluster to obtain a per-slide mean phenotype score vector. Phenotype score distributions were summarised separately for the training, validation, and test splits, stratified by MYCN status. We visualised these distributions using kernel density estimates and violin plots to compare how tiles from each molecular subtype were assigned across the learned GMM phenotypic clusters.

### 4.7 Slide-level phenotype-composition clustering

To test whether slide-level phenotype composition alone grouped tumours by MYCN-associated morphology, we clustered all 189 slides using only their per-slide mean phenotype score vectors; MYCN status, sample type and survival were not used for clustering. The analysis was restricted to the four tumour-associated phenotypic clusters (clusters 2, 3, 5 and 6); cluster 1 (paucicellular tissue of varied composition) and cluster 4 (artefact) were excluded, the former as corresponding to no single tumour-associated tissue state, the latter as reflecting image quality rather than tumour-cell morphology. For each slide, the four scores were renormalised to sum to one, log-transformed (log_10_, with values floored at 10*^−^*^3^), and standardised to zero mean and unit variance across slides so that the dominant tumour clusters (3 and 5) did not swamp the smaller necrotic components. Slides were clustered by Ward’s linkage on Euclidean distances with optimal leaf ordering, and the dendrogram was cut into five slide-level clusters for visualisation and summary in Fig. 3. MYCN status, sample type and survival were added only as post hoc annotations. To quantify MYCN enrichment, the two necrosis-rich slide-level clusters were combined and compared with the remaining three clusters using Fisher’s exact test; the odds ratio and 95% confidence interval were reported as descriptive effect sizes. After clustering, a specialist paediatric pathologist (J.C.H.) reviewed whole-slide images and their tiles from each slide-level cluster and described the tissue on them, including components not captured by the four phenotype scores used for clustering; the per-cluster descriptions in Fig. 3 draw on this review. One representative slide and three representative H&E tiles per cluster were selected from within that cluster in consultation with the pathologist to illustrate the described features. Neither the review nor the selection influenced cluster membership, the five-cluster cut or any statistic derived from the clusters.

### 4.8 Pathology review of learned phenotypic clusters

For downstream interpretability, the selected six-cluster GMM was applied post hoc to GMM-projected tile embeddings across the dataset to compute phenotype scores and hard labels for each cluster. Hard labels were assigned by selecting the cluster with the highest phenotype score for each tile. For each cluster, the ten slides with the highest mean GMM phenotype score for that cluster were identified separately within the MYCN-amplified and non-amplified groups. For all six phenotypic clusters, representative hard-assigned tiles from these slides were retrieved and reviewed by a specialist paediatric pathologist (J.C.H.) to assign descriptive pathology labels. For the two tumour-parenchyma clusters (clusters 3 and 5), representative tiles were additionally reviewed separately by MYCN status. Each cluster was assigned descriptive pathology labels according to the dominant recurrent histopathological patterns observed among the representative tiles and on the whole-slide images from which they were taken. This pathology review was performed after model training and did not influence model fitting, validation, or baseline evaluation.

### 4.9 Cell-level spatial profiling

To characterise the cellular features of MYCN-associated phenotypic clusters, we performed cell-level spatial profiling on tiles from the four MYCN-associated phenotypic clusters (cluster 2, haemorrhagic necrosis; cluster 3, loose tumour; cluster 5, dense tumour; cluster 6, acellular necrosis) (Fig. 4, Fig. 5). For each phenotypic cluster, the ten slides with the highest mean GMM phenotype score for that cluster were identified separately within the MYCN-amplified and non-amplified groups. All tiles whose argmax GMM phenotype score corresponded to that cluster were selected from these slides, yielding 801 tiles for cluster 2 (513 non-amplified, 288 MYCN-amplified), 1,465 for cluster 3 (1,194 non-amplified, 271 MYCN-amplified), 1,574 for cluster 5 (1,245 non-amplified, 329 MYCN-amplified), and 391 for cluster 6 (201 non-amplified, 190 MYCN-amplified). Because necrotic (clusters 2 and 6) tiles contain few segmentable nuclei, fewer cell-level features were available for these two clusters than for the tumour clusters.

Nuclei within selected tiles were segmented and classified using PanNuke-pretrained HoVer-Net [19, 28]. In our neuroblastoma dataset, nuclei labelled as epithelial by HoVer-Net typically corresponded to neuroblasts and were therefore treated as neuroblast cells. The “other” category was excluded from downstream analysis because it frequently included misclassified nuclei, including ganglion cells. The retained cell classes for downstream profiling were neuroblast, immune, and dead cells. Following nuclear segmentation and classification, we extracted handcrafted cell-level features using the SI-MIL framework [29]. Features with more than 50% missing values within either MYCN amplification subgroup were excluded, as were constant-value features. This filtering retained 92 features per tile for downstream latent-space characterisation and statistical modelling. These features captured morphometric, textural, and spatial properties of nuclei and were aggregated at the tile level using summary statistics, including mean, standard deviation, skewness, and kurtosis. Morphometric and textural descriptors included intensity, shape, and texture measurements computed for each retained cell class. Spatial descriptors were computed from nuclei centroids and class labels, including graph-based features derived from cell graphs, as well as entropy- and infiltration-based measures designed to quantify cellular heterogeneity and intermixing among cell types.

### 4.10 Latent space characterisation

To assign each tile a MYCN-amplification score, we trained an L2-regularised logistic regression classifier (regularisation strength *C* = 0.1, class-balanced weights, L-BFGS solver, maximum 1000 iterations) separately for each of the four MYCN-associated clusters (clusters 2, 3, 5, and 6) using the 92 cell-level features per tile described above. Missing feature values were imputed by the per-feature median and all features were standardised to zero mean and unit variance prior to model fitting. The predicted probability of MYCN amplification *p*^ *∈* [0, 1] output by the trained classifier was used as a per-tile soft label, providing a continuous multiparametric characterisation of each tile’s position in the cell-level feature space (Fig. 4 **a, b**). To confirm that the resulting subtype separation did not simply reflect in-sample fitting, we repeated the classification under leave-one-slide-out cross-validation, refitting the median imputer, the standardiser and the classifier on the remaining slides and scoring each held-out slide’s tiles, and summarised the separation by the area under the ROC curve of the per-slide mean out-of-fold soft label.

At the slide level, a per-slide mean feature vector was computed for each cluster by averaging the 92 scaled features across all tiles assigned to a given slide and phenotypic cluster. For each cluster, principal component analysis (PCA) was applied to the resulting 20 *×* 92 matrix of slide-level feature vectors (10 MYCN-amplified and 10 non-amplified slides) to project slides onto a two-dimensional latent space (Fig. 4 **c-f**).

Uniform Manifold Approximation and Projection (UMAP) [30] was applied to the tile-level scaled features of each cluster (*n* = 801, 1, 465, 1, 574, and 391 tiles for clusters 2, 3, 5, and 6, respectively) with parameters *n*_neighbours_ = 15, min dist = 0.1, and a fixed random seed. The resulting two-dimensional embeddings were coloured by the per-tile soft label to visualise the continuous phenotypic gradient across tile sub-populations. Because UMAP preserves local neighbourhood structure rather than global geometry, distances between separated groups in these embeddings are not quantitatively interpretable. The cluster 3 embedding, annotated with representative H&E tiles, is shown in Fig. 4 **g**, and the cluster 5 embedding in Supplementary Fig. B3.

### 4.11 Statistical analysis

Throughout, the slide (tumour sample), rather than the patient, was the unit of analysis, and tiles within a slide were treated as repeated measurements. To test whether the per-tile phenotype-score distributions differed between MYCN-amplified and non-amplified slides (Fig. 2 **a-c**, Supplementary Table A4), we compared, for each phenotypic cluster, the pooled per-tile phenotype scores of the two groups using a two-sample Kolmogorov–Smirnov statistic. Significance was obtained from a permutation null that shuffled MYCN labels at the slide level (2,000 permutations; tiles subsampled to at most 1,500 per slide), preserving the tile-within-slide structure.

To identify features associated with MYCN amplification status while accounting for tile-within-slide correlation, we fitted linear mixed-effects models (LME) separately for each cell-level feature and each phenotypic cluster. Each model included MYCN amplification status as a fixed effect and a random intercept per slide, with the ten slides per subtype (*n* = 10) serving as the statistical units and individual tiles as repeated measurements within slides. *z*-statistics and associated *p*-values were extracted from each model, and Benjamini-Hochberg false discovery rate (FDR) correction was applied across all features per cluster. LME results for all features significant in at least one cluster are provided in Supplementary Table A5.

To attribute the classifier’s predictions to individual features, SHAP (SHapley Additive exPlanations) [31] values were computed using a linear SHAP explainer applied to the logistic regression model described above. The background reference distribution was set to the per-feature training-data mean. SHAP values were computed with respect to the MYCN-amplified class for each tile, and features were ranked by their mean absolute SHAP value across all tiles within each phenotypic cluster. Results are shown as beeswarm plots in Fig. 5 **b, c** and Supplementary Fig. B4.

### 4.12 Survival analysis

Overall survival was analysed at the slide level. Survival time started at the date of the biopsy or resection from which the FFPE block was taken. For patients who had died, it ended at the recorded date of death. Patients recorded as alive were censored at a common study cut-off, set to the latest date of death observed in the cohort (9 March 2024). Survival times were expressed in months (days divided by 30.44). Of the 189 slides in the model cohort, 187 were analysed; two slides from a single patient were excluded because the recorded date of death preceded the resection date, which yields a negative, non-interpretable survival time. Because the unit of analysis was the slide and a patient could contribute more than one slide, within-patient correlation was not modelled, and all log-rank p-values are reported as descriptive. For the phenotype-composition clusters in Fig. 3, Kaplan-Meier curves were drawn descriptively for each of the five slide-level clusters; the two necrosis-rich clusters were also combined and compared with the remaining three clusters by an unadjusted log-rank test.

Kaplan–Meier curves were estimated by per-slide FISH MYCN status (Fig. 6 **a**). A per-slide predicted probability of MYCN amplification was obtained out-of-fold, by applying each stratified hold-out split’s trained Pheno-MYCN model to the slides it had held out for testing and averaging the probabilities for any slide held out in more than one split. A binary H&E MYCN call was then assigned at a threshold of 0.5. This analysis was restricted to the 79 slides that received such an out-of-fold prediction. Kaplan–Meier curves were estimated by the predicted call and compared with the FISH benchmark on the same slides (Fig. 6 **b**). The continuous predicted probability was additionally evaluated in a Cox proportional-hazards model (hazard ratio per standard deviation) and summarised by Harrell’s concordance index, which was compared with that of a Cox model fitted to binary FISH status on the same slides.

To test whether the morphology adds prognostic information to the molecular assay, three Cox proportional-hazards models were fitted on the 79 slides with an out-of-fold prediction: FISH MYCN status alone, the continuous H&E-derived MYCN score alone (standardised to zero mean and unit variance), and the two together. Each model was summarised by Harrell’s concordance index computed out of fold, to avoid in-sample optimism: slides were divided into five folds, the model was refitted on four folds and used to predict the log partial hazard of the held-out fold, and a single concordance index was computed on the pooled out-of-fold predictions. This was repeated for 20 random fold assignments, and the mean and standard deviation across repeats are reported (Fig. 6 **d**). The contribution of the morphology score over and above FISH status was assessed by a likelihood-ratio test comparing the FISH-only and FISH-plus-morphology models (one degree of freedom). A combined risk score was then defined as the out-of-fold linear predictor of the FISH-plus-morphology model, split at the cohort median, and the resulting low- and high-risk groups were compared by Kaplan-Meier estimation and the log-rank test (Fig. 6 **c**).

To ask whether the phenotypic clusters themselves carry prognostic information, each slide was summarised by two statistics for every cluster: the mean phenotype score, the average of the tile-level posterior phenotype scores for that cluster across the slide, and the dominant-tile fraction, the proportion of the slide’s tiles whose phenotypic score for that cluster exceeds 0.5. Each score was standardised to zero mean and unit variance and analysed by Cox proportional-hazards models, with the hazard ratio reported per standard deviation, fitted both univariably and after adding FISH MYCN status as a covariate (Supplementary Table A7). The clusters were then tested jointly rather than one at a time. The four tumour-associated clusters (2, 3, 5 and 6) were expressed as a compositional profile using the same transform as the slide-level clustering, renormalised to sum to one per slide, log-transformed (log_10_, floored at 10*^−^*^3^) and standardised across slides, and were entered together into a Cox model. Composition-only, MYCN-only and MYCN-plus-composition models were compared by the same repeated out-of-fold concordance index described above, and the contribution of the composition over and above MYCN status was assessed by a likelihood-ratio test (four degrees of freedom).

## Data availability

The whole-slide images and linked clinical data analysed in this study were provided by the Great Ormond Street Hospital biobank and were collected under the Stratified Medicine Paediatrics (SMPaeds) and Stratified Medicine Paediatrics 2 (SMPaeds2) programmes, sponsored by The Institute of Cancer Research. These data are available under restricted access because whole-slide images and the associated clinical records are potentially patient-identifiable, and their release is governed by the terms of the studies’ ethical approvals and the associated data-sharing agreements. Requests for access should be directed to The Institute of Cancer Research and will be assessed by the Stratified Medicine Paediatrics Study Management Group against the studies’ data access policies. Access, where granted, requires a data use agreement restricting use to non-commercial academic research and prohibiting onward redistribution and any attempt at re-identification, and is conditional on appropriate institutional and ethical approvals. No patient-identifiable information is released.

## Code availability

The implementation of Pheno-MYCN, comprising the model, training and inference pipelines and the analysis scripts used to generate all figures and tables, is publicly available at: https://github.com/cbhindex/pheno mycn under a GPL-3.0 licence. The trained model weights are released for non-commercial academic research use only, consistent with the terms of the UNI encoder [16] on whose embeddings the model was trained. Analyses were run under Python 3.10 with PyTorch 2.0.1 (CUDA 11.8); the complete tested environment is specified in environment.yml and requirements.txt in the repository.

## Author contribution statement

B.C. and O.F. contributed equally to this work; C.B. is the lead contact. B.C.: Data curation, Formal analysis, Investigation, Methodology, Software, Validation, Visualization, Writing – original draft, Writing – review & editing. O.F.: Conceptualization, Formal analysis, Investigation, Methodology, Software, Validation, Visualization, Writing – original draft, Writing – review & editing. R.N.: Data curation, Investigation, Methodology, Writing – review & editing. M.D.V.: Investigation, Methodology. S.W.A.: Data curation, Resources. T.J.: Investigation, Resources. S.G.: Funding acquisition, Resources, Writing – review & editing. L.C.: Funding acquisition, Resources, Writing – review & editing. J.C.H.: Investigation, Resources, Validation, Visualization, Writing – review & editing. C.B.: Conceptualization, Funding acquisition, Project administration, Resources, Supervision, Validation, Writing – review & editing.

## Acknowledgments

This work was funded by Cancer Research UK and Children with Cancer UK through Stratified Medicine Paediatrics (SMPaeds), under grant number CRCEMA-Dec21/100003, and Stratified Medicine Paediatrics 2 (SMPaeds2), under grant number CRCEMA-Jul23/100001. We thank Great Ormond Street Hospital Biobank for the provision of human tissue samples and clinical data, and the patients and their families who contributed samples and data to this study. We are grateful to the Institute of Cancer Research for research support, and to all healthcare workers who cared for the patients without whose input this work would not have been possible.

## Appendix A Supplementary Tables

**Table A1:** Cohort demographics stratified by MYCN amplification status. Columns group patients by MYCN status (amplified at any timepoint); slide rows count the slides contributed by each patient group.

|  | *MYCN* non-amplified<br>(*n* = 58) | *MYCN* amplified<br>(*n* = 28) | Overall<br>(*n* = 86) |
| --- | --- | --- | --- |
| Patients, *n* | 58 | 28 | 86 |
| Age at biopsy, *y* — median (IQR) | 6.6 (4.6–9.0) | 4.3 (2.5–5.8) | 5.7 (3.8–8.6) |
| Age < 18 months | 1 (1.7) | 2 (7.1) | 3 (3.5) |
| Age 18 months – 5 years | 18 (31.0) | 15 (53.6) | 33 (38.4) |
| Age ≥ 5 years | 39 (67.2) | 11 (39.3) | 50 (58.1) |
| Sex |  |  |  |
| Female | 32 (55.2) | 12 (42.9) | 44 (51.2) |
| Male | 26 (44.8) | 16 (57.1) | 42 (48.8) |
| Patient status |  |  |  |
| Patient deceased | 35 (60.3) | 25 (89.3) | 60 (69.8) |
| Patient alive | 23 (39.7) | 3 (10.7) | 26 (30.2) |
| Slides, *n* | 130 | 59 | 189 |
| Timepoint |  |  |  |
| Primary | 46 (35.4) | 22 (37.3) | 68 (36.0) |
| Relapse | 84 (64.6) | 37 (62.7) | 121 (64.0) |

Data are *n* (%) unless stated; percentages exclude unknowns from the denominator. Patient *MYCN* status is positive if any slide is amplified. Slide rows count the slides contributed by each patient group (130/59 = 189); by per-slide label the split is 140 non-amplified / 49 amplified, because the 28 amplified patients contributed 49 amplified and 10 non-amplified slides.

**Table A2:** GMM phenotypic cluster number selection: mean classification performance (*±* SD) across ten stratified hold-out splits for Pheno-MYCN trained with *K* = 2 to 9 GMM phenotypic clusters. **Bold** indicates the selected configuration. *^†^K* = 4 yields a marginally higher mean AUC (0.917) but lower accuracy (0.856) and F1-score (0.813).

| $K$ | Accuracy | Precision | Recall | F1-score | AUC |
| --- | --- | --- | --- | --- | --- |
| 2 | $0.822 \pm 0.141$ | $0.847 \pm 0.152$ | $0.775 \pm 0.152$ | $0.783 \pm 0.160$ | $0.911 \pm 0.134$ |
| 3 | $0.800 \pm 0.172$ | $0.809 \pm 0.194$ | $0.758 \pm 0.182$ | $0.766 \pm 0.187$ | $0.861 \pm 0.155$ |
| 4<sup>†</sup> | $0.856 \pm 0.105$ | $0.899 \pm 0.099$ | $0.800 \pm 0.131$ | $0.813 \pm 0.133$ | $0.917 \pm 0.106$ |
| 5 | $0.822 \pm 0.130$ | $0.844 \pm 0.150$ | $0.775 \pm 0.147$ | $0.783 \pm 0.153$ | $0.883 \pm 0.106$ |
| **6** | **$0.889 \pm 0.117$** | **$0.894 \pm 0.129$** | **$0.867 \pm 0.125$** | **$0.873 \pm 0.127$** | **$0.900 \pm 0.122$** |
| 7 | $0.789 \pm 0.110$ | $0.818 \pm 0.119$ | $0.750 \pm 0.118$ | $0.747 \pm 0.121$ | $0.883 \pm 0.115$ |
| 8 | $0.844 \pm 0.078$ | $0.863 \pm 0.105$ | $0.800 \pm 0.105$ | $0.809 \pm 0.103$ | $0.889 \pm 0.094$ |
| 9 | $0.822 \pm 0.130$ | $0.840 \pm 0.146$ | $0.783 \pm 0.137$ | $0.789 \pm 0.144$ | $0.872 \pm 0.091$ |

**Table A3:** Per-split test set performance for Pheno-MYCN and baseline models across all ten stratified hold-out splits. The mean *±* SD row for each model corresponds to the values reported in Table 1 of the main text.

| Fold | Accuracy | Precision | Recall | F1-score | AUC |
| --- | --- | --- | --- | --- | --- |
| *Pheno-MYCN* |  |  |  |  |  |
| 0 | 0.667 | 0.650 | 0.667 | 0.649 | 0.833 |
| 1 | 0.889 | 0.929 | 0.833 | 0.862 | 0.944 |
| 2 | 1.000 | 1.000 | 1.000 | 1.000 | 1.000 |
| 3 | 0.889 | 0.929 | 0.833 | 0.862 | 0.667 |
| 4 | 0.889 | 0.929 | 0.833 | 0.862 | 0.889 |
| 5 | 0.778 | 0.750 | 0.750 | 0.750 | 0.944 |
| 6 | 1.000 | 1.000 | 1.000 | 1.000 | 1.000 |
| 7 | 1.000 | 1.000 | 1.000 | 1.000 | 1.000 |
| 8 | 1.000 | 1.000 | 1.000 | 1.000 | 1.000 |
| 9 | 0.778 | 0.750 | 0.750 | 0.750 | 0.722 |
| **Mean ± SD** | **0.889 ± 0.117** | **0.894 ± 0.129** | **0.867 ± 0.125** | **0.873 ± 0.127** | **0.900 ± 0.122** |
| *CLAM-SB* |  |  |  |  |  |
| 0 | 0.889 | 0.929 | 0.833 | 0.862 | 0.889 |
| 1 | 0.778 | 0.875 | 0.667 | 0.679 | 0.778 |
| 2 | 1.000 | 1.000 | 1.000 | 1.000 | 1.000 |
| 3 | 0.667 | 0.333 | 0.500 | 0.400 | 0.833 |
| 4 | 0.778 | 0.875 | 0.667 | 0.679 | 0.778 |
| 5 | 0.889 | 0.929 | 0.833 | 0.862 | 0.778 |
| 6 | 0.778 | 0.875 | 0.667 | 0.679 | 0.833 |
| 7 | 0.778 | 0.875 | 0.667 | 0.679 | 0.667 |
| 8 | 0.889 | 0.929 | 0.833 | 0.862 | 0.944 |
| 9 | 0.778 | 0.750 | 0.750 | 0.750 | 0.667 |
| **Mean ± SD** | **0.822 ± 0.094** | **0.837 ± 0.188** | **0.742 ± 0.139** | **0.745 ± 0.164** | **0.817 ± 0.108** |
| *TransMIL* |  |  |  |  |  |
| 0 | 0.778 | 0.750 | 0.750 | 0.750 | 0.889 |
| 1 | 0.778 | 0.875 | 0.667 | 0.679 | 0.778 |
| 2 | 1.000 | 1.000 | 1.000 | 1.000 | 1.000 |
| 3 | 0.889 | 0.929 | 0.833 | 0.862 | 0.833 |
| 4 | 0.889 | 0.929 | 0.833 | 0.862 | 0.889 |
| 5 | 0.889 | 0.929 | 0.833 | 0.862 | 0.889 |
| 6 | 0.667 | 0.333 | 0.500 | 0.400 | 0.611 |
| 7 | 0.778 | 0.875 | 0.667 | 0.679 | 0.944 |
| 8 | 0.778 | 0.750 | 0.750 | 0.750 | 0.889 |
| 9 | 0.778 | 0.750 | 0.750 | 0.750 | 0.722 |
| **Mean ± SD** | **0.822 ± 0.094** | **0.812 ± 0.190** | **0.758 ± 0.133** | **0.759 ± 0.160** | **0.844 ± 0.114** |

**Table A4:**
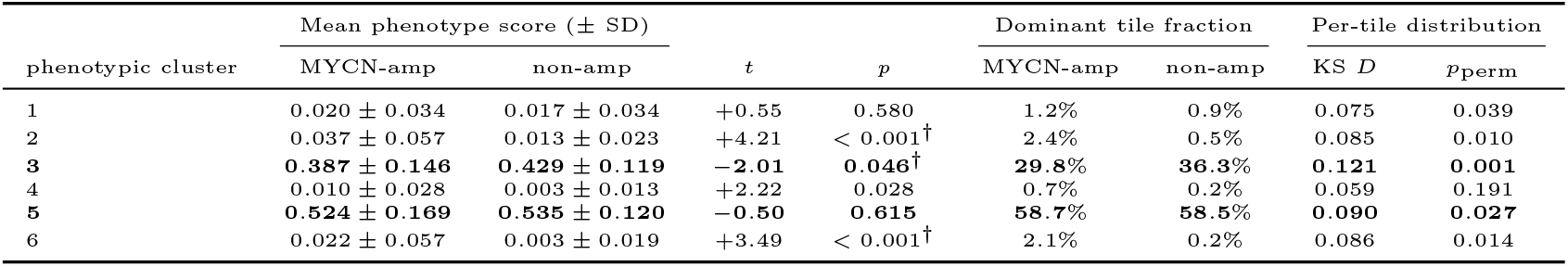
Slide-level GMM phenotype score statistics across all 189 slides (MYCN-amplified *n* = 49; non-amplified *n* = 140). Mean phenotype score is the per-slide average of tile-level posterior probabilities for each GMM phenotypic cluster; values are reported as mean *±* SD across slides. The dominant tile fraction is the proportion of tiles per slide for which a given phenotypic cluster has a posterior probability *>* 0.5 averaged across slides. Two-sample *t*-tests compare the mean phenotype score between MYCN-amplified and non-amplified slides. The per-tile distribution columns report a two-sample Kolmogorov–Smirnov statistic (*D*) comparing the pooled per-tile phenotype-score distributions of MYCN-amplified versus non-amplified slides, with *p*_perm_ obtained from 2000 slide-label permutations (tiles capped at 1500 per slide), which preserve the tile-within-slide structure. **Bold** rows indicate the dominant tumour phenotypic clusters (3 and 5) examined in main-text detail; all six phenotypic clusters were reviewed by the pathologist, and the four MYCN-associated clusters (2, 3, 5 and 6) were carried forward to cell-level profiling, with clusters 2 and 6 detailed in the Supplementary Figures. *^†^* Direction of enrichment was consistent across the validation and test splits (see main text). All reported *p* values are nominal and uncorrected for multiple comparisons across the six phenotypic clusters; under a conservative Bonferroni threshold (*α* = 0.05*/*6 *≈* 0.008), the slide-level enrichment of clusters 2 and 6 (*t*-test) and the cluster-3 per-tile distribution difference (KS) remain significant, while the remaining nominally significant values (*p <* 0.05) should be interpreted with caution.

**Table A5:** Linear mixed-effects model (LME) results for cell-level features in clusters 2, 3, 5 and 6. Features with *p*_(BH-adjusted)_ *<* 0.05 in at least one phenotypic cluster are shown, grouped by feature category and ordered by effect magnitude. *z*: fixed-effect *z*-statistic; negative values indicate reduction in MYCN-amplified tiles. *p*_(BH-adjusted)_: Benjamini–Hochberg adjusted *p*-value (*^∗∗∗^p <* 0.001; *^∗∗^p <* 0.01; *^∗^p <* 0.05; –, absent from the cluster’s set).

| Modality | Feature | Mixed-effects $z$-statistic | | | |
| --- | --- | --- | --- | --- | --- |
|  |  | C2 | C3 | C5 | C6 |
| **Connectivity** | mean of cells’ clustering_coefficient | −1.8 | −5.7*** | −2.5 | – |
|  | max of cells’ closeness centrality | −1.9 | −5.7*** | −3.5** | – |
|  | max of cells’ clustering_coefficient | −1.9 | −5.6*** | −2.0 | – |
|  | mean of cells’ closeness centrality | −1.8 | −5.6*** | −3.5** | – |
|  | min of cells’ closeness centrality | −1.7 | −5.6*** | −3.5** | – |
|  | max of cells’ degree | −2.2 | −5.5*** | −1.9 | – |
|  | mean of cells’ degree | −2.0 | −5.4*** | −1.8 | – |
|  | min of cells’ degree | −2.0 | −5.4*** | −1.7 | – |
|  | min of cells’ clustering_coefficient | −1.7 | −5.4*** | −2.8* | – |
|  | max of cells’ degree centrality | −1.9 | −5.2*** | −3.8** | – |
|  | std of cells’ clustering_coefficient | −2.2 | −5.1*** | −1.2 | – |
|  | std of cells’ degree | −2.3 | −5.1*** | −2.4 | – |
|  | std of cells’ closeness centrality | −2.2 | −5.1*** | −3.1* | – |
|  | mean of cells’ degree centrality | −1.7 | −4.9*** | −3.7** | – |
|  | min of cells’ degree centrality | −1.6 | −4.9*** | −3.6** | – |
|  | std of cells’ degree centrality | −2.2 | −4.7*** | −3.8** | – |
|  | skew of cells’ degree centrality | −1.5 | −4.0*** | −0.5 | – |
|  | skew of cells’ degree | −1.5 | −4.0*** | −0.5 | – |
|  | skew of cells’ closeness centrality | −1.7 | −3.9*** | −0.8 | – |
|  | kurtosis of cells’ clustering_coefficient | +2.2 | +3.4** | +2.1 | – |
| **Diversity** | Global Simpson index | −2.1 | −4.7*** | −0.7 | – |
|  | Local richness skew | −2.2 | −4.4*** | −0.7 | – |
|  | Global Shannon index | −1.0 | −4.4*** | −2.0 | – |
|  | Global Richness (number of cell-types present) | −1.4 | −4.3*** | −0.4 | – |
|  | Local Renyi entropy (at alpha=infinity) skew | −2.0 | −4.1*** | −0.4 | – |
|  | Global Renyi entropy (at alpha=infinity) | −1.3 | −4.0*** | −2.0 | – |
|  | Local Shannon index skew | −2.1 | −3.0** | −0.6 | – |
| **Morphology** | kurtosis of their orientation | – | −3.7*** | −3.6** | – |
|  | skew of their axis_minor.length | – | +2.5* | +1.0 | – |
| **Colour/texture** | mean_h | +5.2*** | – | – | +3.6** |
|  | mean_r | +3.9** | – | – | +2.8 |
|  | skew of their intensity_mean | – | −3.4** | −2.0 | – |
|  | mean_v | +3.4* | – | – | +2.6 |
| **Counts** | number of necrosis cells | −0.1 | +2.7* | +3.0* | −0.5 |

**Table A6:** Top ten features by mean absolute SHAP value for each MYCN-associated phenotypic cluster. SHAP values were computed with a linear explainer on the L2-regularised logistic-regression classifier of tile-level MYCN-amplification status. Features are ranked within each cluster by mean *|*SHAP*|*. The *Direction* column indicates the class a feature predominantly drives the prediction toward: “MYCN-amp” when the mean SHAP value across MYCN-amplified tiles is positive and “non-amp” otherwise. Colour/texture channels named as in Table A5 (`glcm` = grey-level co-occurrence texture; `mean_gray` = greyscale intensity).

| Feature | Mean $\vert \text{SHAP}\vert $ | Direction |
| --- | --- | --- |
| **Cluster 2 (haemorrhagic necrosis)** |  |  |
| std_b | 0.697 | MYCN-amp |
| mean_h | 0.410 | MYCN-amp |
| mean_r | 0.215 | MYCN-amp |
| std_s | 0.210 | MYCN-amp |
| glcm_homogeneity | 0.187 | non-amp |
| std_r | 0.175 | MYCN-amp |
| std_v | 0.164 | non-amp |
| mean_gray | 0.160 | MYCN-amp |
| area | 0.159 | non-amp |
| mean_g | 0.120 | non-amp |
| **Cluster 3 (loose tumour)** |  |  |
| Local Simpson index skew | 0.467 | MYCN-amp |
| number of neuroblast cells | 0.399 | MYCN-amp |
| std of their perimeter | 0.303 | MYCN-amp |
| Local richness skew | 0.302 | MYCN-amp |
| std of their area | 0.283 | non-amp |
| max of cells' clustering_coefficient | 0.261 | MYCN-amp |
| Global Simpson index | 0.246 | MYCN-amp |
| Global Shannon index | 0.240 | MYCN-amp |
| std of cells' degree centrality | 0.237 | non-amp |
| skew of their area | 0.228 | MYCN-amp |
| **Cluster 5 (dense tumour)** |  |  |
| Global Shannon index | 0.320 | MYCN-amp |
| std of their perimeter | 0.319 | MYCN-amp |
| Global Richness (number of cell-types present) | 0.312 | MYCN-amp |
| perimeter | 0.254 | non-amp |
| eccentricity | 0.251 | MYCN-amp |
| Local richness skew | 0.231 | MYCN-amp |
| std of their axis_major_length | 0.218 | non-amp |
| kurtosis of their perimeter | 0.211 | non-amp |
| number of necrosis cells | 0.182 | MYCN-amp |
| Local Renyi entropy (at alpha=infinity) skew | 0.175 | MYCN-amp |
| **Cluster 6 (acellular necrosis)** |  |  |
| std_b | 0.429 | MYCN-amp |
| mean_h | 0.329 | MYCN-amp |
| mean_g | 0.277 | non-amp |
| mean_gray | 0.234 | non-amp |
| mean_s | 0.226 | non-amp |
| std_v | 0.164 | MYCN-amp |
| mean_b | 0.162 | MYCN-amp |
| tissue_area_fraction | 0.161 | MYCN-amp |
| mean_r | 0.151 | MYCN-amp |
| std_s | 0.137 | MYCN-amp |

**Table A7:** Cox proportional-hazards models for the per-slide phenotype-cluster scores, each cluster entered on its own, fitted univariably and with adjustment for FISH MYCN status (slide-level cohort, *n* = 187). Hazard ratios are per one standard deviation of the score, with 95% confidence intervals. The adverse univariable associations (C2 and C6) lose significance after adjustment for MYCN, so no single cluster is prognostic independent of MYCN status. Because entering clusters one at a time cannot capture their joint structure, the four tumour-associated clusters were additionally entered together as a compositional profile and evaluated out of fold against MYCN status; that analysis is reported in the main text and likewise found no independent prognostic signal.

| Cluster | Univariable |  | Adjusted for MYCN |  |
| --- | --- | --- | --- | --- |
| | HR (95% CI) | $p$ | HR (95% CI) | $p$ |
| *Dominant-tile fraction* |  |  |  |  |
| C2 | 1.23 (1.05–1.43) | 0.009 | 1.14 (0.97–1.34) | 0.120 |
| C3 | 0.92 (0.77–1.09) | 0.338 | 0.95 (0.81–1.13) | 0.587 |
| C5 | 1.01 (0.85–1.20) | 0.937 | 1.02 (0.86–1.21) | 0.821 |
| C6 | 1.19 (1.02–1.39) | 0.025 | 1.11 (0.93–1.31) | 0.245 |
| *Mean phenotype score* |  |  |  |  |
| C2 | 1.24 (1.06–1.45) | 0.007 | 1.14 (0.96–1.34) | 0.129 |
| C3 | 0.91 (0.76–1.09) | 0.303 | 0.95 (0.80–1.13) | 0.562 |
| C5 | 0.99 (0.82–1.18) | 0.874 | 1.01 (0.85–1.20) | 0.891 |
| C6 | 1.19 (1.03–1.39) | 0.021 | 1.11 (0.94–1.31) | 0.232 |

## Appendix B Supplementary Figures

**Fig. B1:**
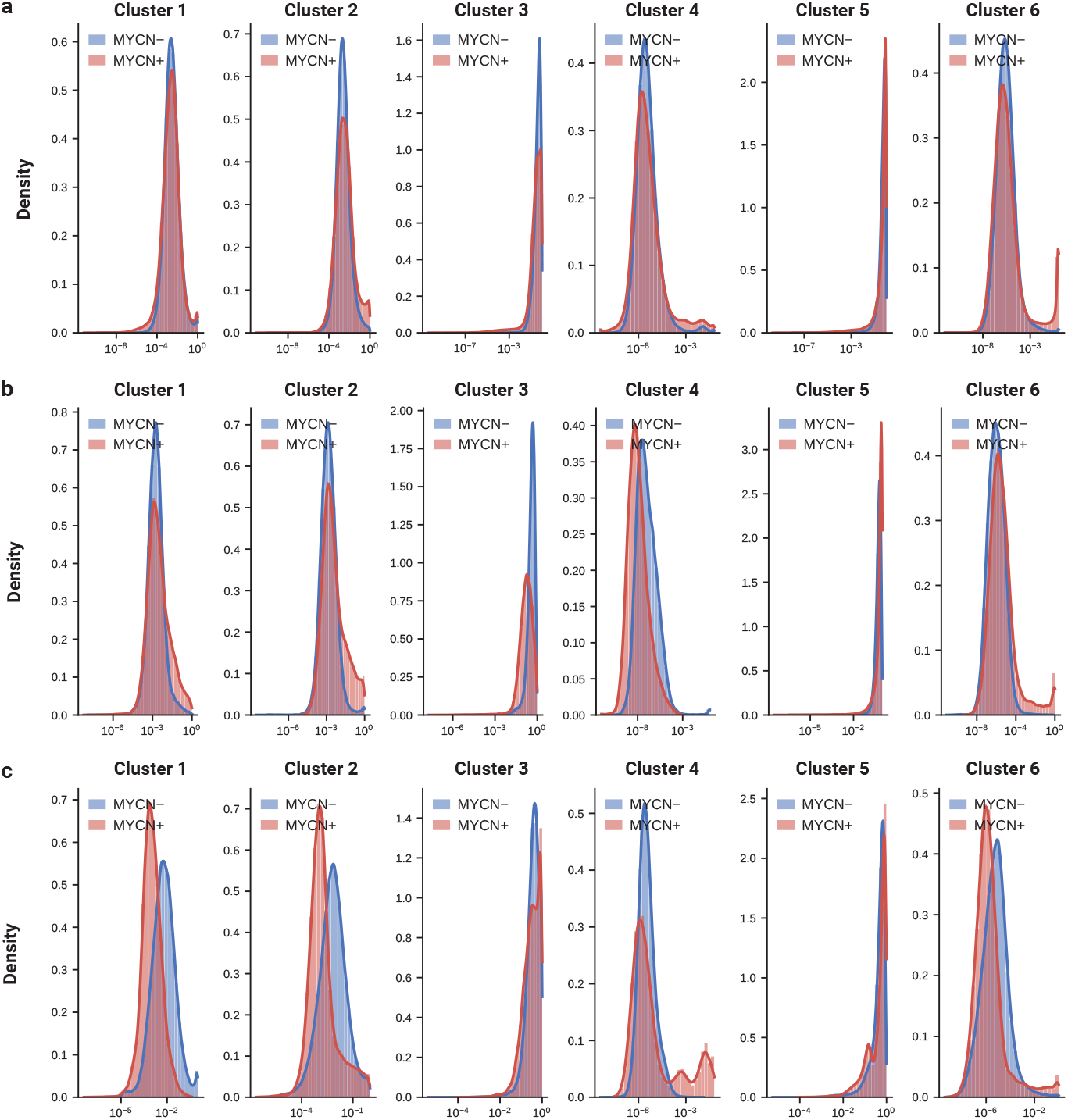
**a, b, c**, Kernel density estimates of per-tile GMM phenotype scores for MYCN-amplified and non-amplified cases across the training (**a**), validation (**b**), and test (**c**) sets.

**Fig. B2:**
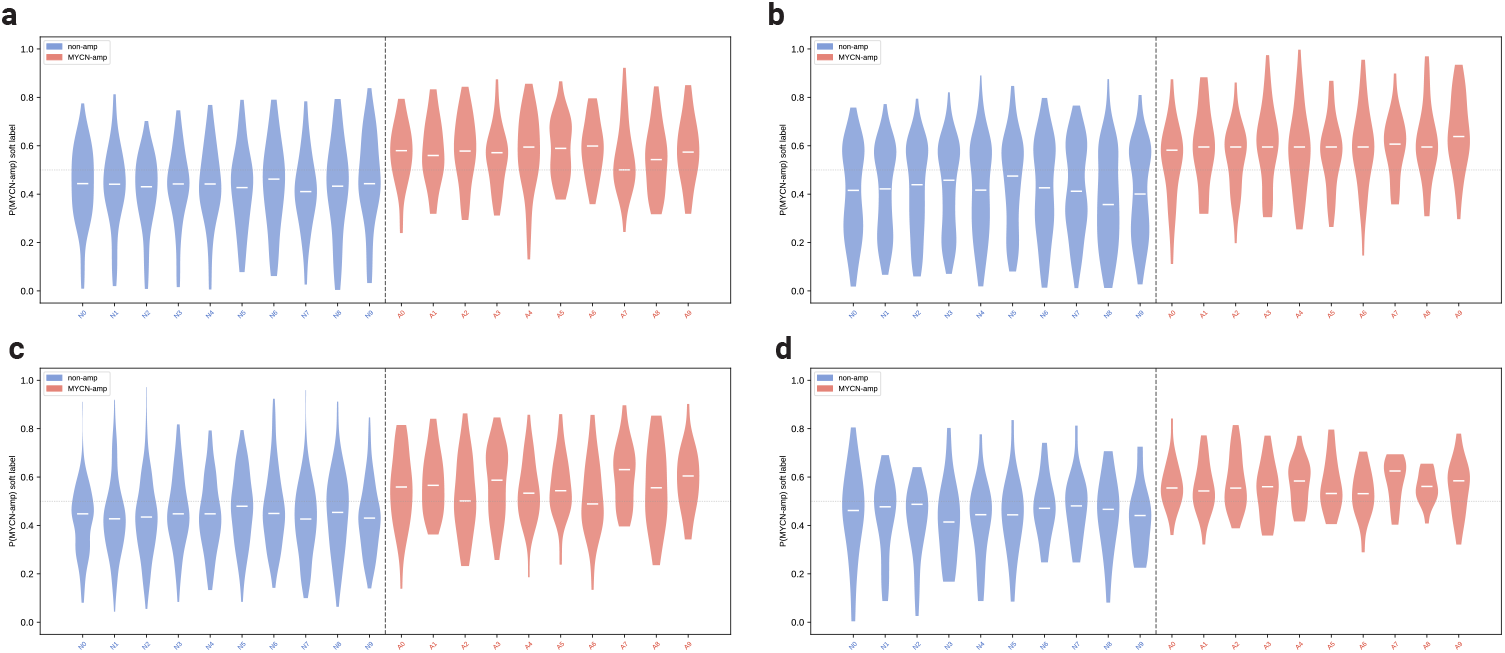
Per-slide distributions of the cell-level soft label for each MYCN-associated phenotypic cluster. Companion to Fig. 4 **a, b**. For each cluster, the distribution of the per-tile soft label is shown separately for every slide (cluster 2, haemorrhagic necrosis, **a**; cluster 3, loose tumour, **b**; cluster 5, dense tumour, **c**; and cluster 6, acellular necrosis, **d**). The soft label is the probability that a tile is MYCN-amplified, from a logistic-regression classifier trained on segmented-cell features (Fig. 4). Within each panel, the dashed vertical line separates the 10 non-amplified slides (blue, N0 to N9; left) from the 10 MYCN-amplified slides (red, A0 to A9; right); each violin pools all tiles of one slide, the white bar marks that slide’s median, and the dotted horizontal line marks 0.5. In every cluster, MYCN-amplified slides sit higher than non-amplified slides, and the shift holds slide-by-slide rather than being driven by a few samples; the spread within each violin reflects intra-tumour (within-slide) heterogeneity. These per-slide distributions are what the main figure summarises as per-slide medians (Fig. 4 **a**) and pooled per-tile densities (Fig. 4 **b**).

**Fig. B3:**
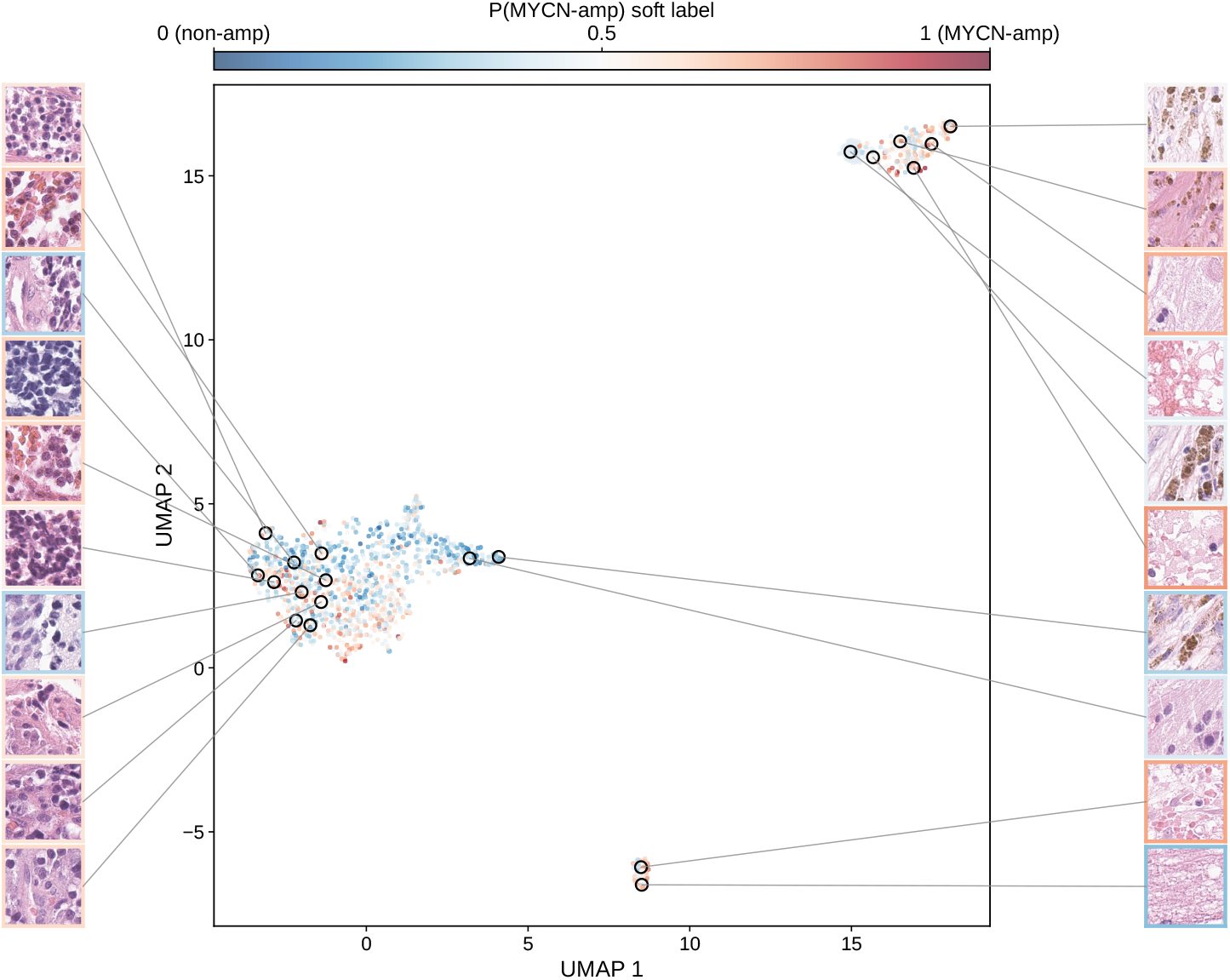
Cell-level latent space of phenotypic cluster 5 (dense tumour): soft-label UMAP with representative tiles. Supplementary companion to Fig. 4 **g**. UMAP embedding of the cluster 5 tiles (one point per tile), coloured by the per-tile soft label; the soft label is the probability that a tile is MYCN-amplified, from a logistic-regression classifier trained on segmented-cell features aggregated per tile (Fig. 4). Representative tiles sampled from the left and right of the embedding are shown around it; each tile’s coloured border encodes its own soft label on the same blue-red scale.

**Fig. B4:**
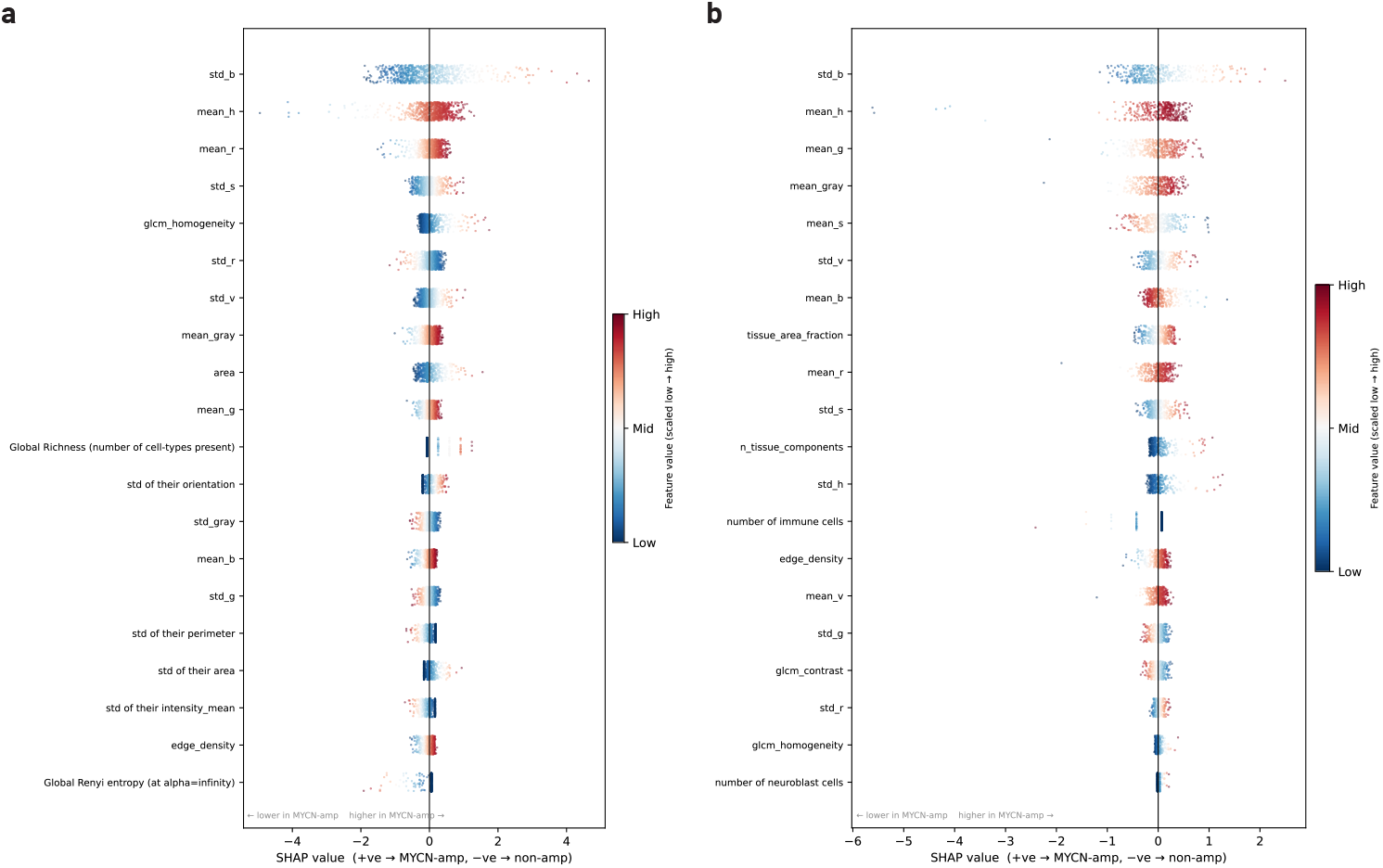
SHAP feature attributions for the haemorrhagic-necrosis (cluster 2, **a**) and acellular-necrosis (cluster 6, **b**) clusters. Each point is one tile; the x-axis is the SHAP value and point colour encodes the scaled feature value. Features are ordered top-to-bottom by mean absolute SHAP value. For both necrotic clusters, the MYCN signal is carried predominantly by colour and texture features rather than the spatial-connectivity and population-diversity features that dominate the tumour clusters. This is consistent with the mixed-effects analysis (Fig. 5 **a**) and with necrosis and haemorrhage being colour-defined, largely acellular tissues. SHAP, SHapley Additive exPlanations.

